# Cellular Maturation of Oligodendrocytes is Governed by Transient Gene Melting

**DOI:** 10.1101/2022.11.17.516981

**Authors:** Kevin C. Allan, Tyler E. Miller, Andrew R. Morton, Marissa A. Scavuzzo, Matthew S. Elitt, Benjamin L.L. Clayton, Lucille R. Hu, Jost K. Vrabic, Hannah E. Olsen, Daniel C. Factor, Jonathan E. Henninger, Richard A. Young, Charles Y. Lin, Peter C. Scacheri, Paul J. Tesar

**Affiliations:** Department of Genetics and Genome Sciences, Case Western Reserve University School of Medicine, Cleveland, Ohio 44106, USA; Department of Molecular and Human Genetics, Baylor College of Medicine, Houston, TX 77030, USA; Whitehead Institute for Biomedical Research, Cambridge, MA 02142, USA; Department of Pathology, Massachusetts General Hospital, Boston, MA 02114, USA

## Abstract

Pluripotent stem cells (PSCs) provide an unlimited source for generating somatic cell types. However, generating fully mature cells constitutes a bottleneck for realizing their full potential in research and medicine. Here, we report a transcriptional mechanism that governs the timing of cellular maturation in post-mitotic oligodendrocytes. During differentiation of PSCs to oligodendrocytes, the transcription factor SOX6 redistributes from nearly all super enhancers in proliferating oligodendrocyte progenitor cells to cluster across specific gene bodies in immature oligodendrocytes. These sites exhibit ‘gene melting’, a process of extensive chromatin decondensation and transcription, which abruptly turns off upon maturation. Suppression of SOX6 deactivates these immaturity loci, resulting in rapid transition to mature myelinating oligodendrocytes. Cells harboring this immature oligodendrocyte SOX6 gene signature are specifically enriched in multiple sclerosis patient brains, suggestive that failed maturation may contribute to limited myelin regeneration in disease. Collectively, our finding that maturation rate is controlled by transient transcriptional clusters may inform approaches to accelerate the generation and regeneration of mature cell types.

**HIGHLIGHTS:**

- Transcription factors (TFs) can act as gatekeepers of post-mitotic cellular maturation
- In oligodendrocytes, clustering of SOX6 controls the immaturity program through transient gene melting
- Suppressing SOX6 deactivates immaturity genes and unlocks oligodendrocyte maturation
- SOX6 immaturity signature is enriched in oligodendrocytes in multiple sclerosis patients

## INTRODUCTION

The potential of stem cells to generate all cell types in the body offers nearly limitless applications for regenerative medicine, disease modeling, and drug discovery.^1–4^ Yet, and despite decades of research efforts, most cellular platforms remain restricted to generating early differentiated cell states that are closer to fetal cells and far removed from mature and fully functional cells in adult tissues.^1–3,5–8^ Increasingly, it has come to be recognized that cell differentiation is but the first step in the process of generating mature tissues. In essence, proliferating progenitor cells are typically specified to their initial post-mitotic state through rapid and global changes in chromatin and transcription. However, within the lifetime of a multicellular organism, the majority of cells exist in an extended postnatal, post-mitotic maturation continuum through which they acquire specialization and functionality by largely unknown processes.^1^ Hence, delineating and manipulating such maturation regulators represents a new frontier in regenerative medicine.

Maturation following differentiation of proliferating progenitor cells is a feature common to multiple cell lineages. In the developing heart, proliferative progenitors first acquire a cardiac fate through differentiation. They subsequently elongate and undergo dramatic alteration in metabolism to become functionally mature cardiomyocytes, with the electrical and contractile properties required for heart function.^9^ Similarly, in the pancreas, developing beta cells must undergo metabolic and genetic adaptations to eventually execute the mature function of insulin release in response to increased glucose levels.^10,11^ To a certain extent, differences in morphology and functionality acquired during cellular maturation can be partly attributed to alterations in transcriptional profiles, as these blueprints provide unique instructions to establish cellular identity and maintain homeostasis.^1,12,13^ Furthermore, delineating master network regulators of cell state transitions in specific lineages can be elucidated by combining transcriptional analysis with chromatin state profiling.^14–17^ However, these approaches have yet to be successfully used to define the controllers of cellular maturation. As a result, the stem cell field is in need of experimental strategies that can rapidly and efficiently generate mature cells for pluripotent stem cell-derived disease modeling, drug screening, and tissue transplantation.

In the central nervous system (CNS), oligodendrocytes first undergo differentiation during fetal development and then, after birth and into adulthood, enter a prolonged maturation process to become mature myelinating oligodendrocytes. Mature oligodendrocytes provide trophic support to neurons and are capable of wrapping neuronal axons in myelin. This feature is unique to vertebrates and allows for rapid propagation of action potentials.^17^ It is also continuous, as oligodendrocyte progenitor cells (OPCs) give rise to immature and highly arborized oligodendrocytes, which then develop into mature oligodendrocytes in the brain and spinal cord throughout life.^18–20^ Understanding trajectories of oligodendrocyte differentiation and subsequent maturation is important because lack of mature oligodendrocyte regeneration drives dysfunction in patients with numerous neurological diseases.^21–24^ Therefore, oligodendrocyte development can be leveraged as a model system to interrogate transcriptional mechanisms governing differentiation and long-term maturation.^18^ Understanding such mechanisms at a cellular and molecular level offers the ability to develop targeted therapies to promote regeneration of myelin in numerous neurological diseases. More broadly, understanding how to generate functionally mature cells has tremendous potential in regenerative medicine applications and disease modeling paradigms.

## RESULTS

### SOX6 is a key regulator of immature oligodendrocyte formation

Leveraging our *in vitro* pluripotent stem cell-derived oligodendrocyte platform, we interrogated the transcriptional and epigenetic landscape of defined states spanning the lineage, including OPCs and immature oligodendrocytes, with the goal of identifying potential gatekeepers of terminal lineage maturation (Figure 1A, S1A, and S1B).^25^ Gene ontology (GO) analysis of differentially expressed genes between *in vitro* OPCs and immature oligodendrocytes (log2FC > 2, P-adj <0.001) and comparison of the transcriptional state of *in vitro* immature oligodendrocytes with cells isolated *in vivo* from the mouse CNS confirm that our culture system generates a highly enriched population of immature, newly specified oligodendrocytes (Figures S1C–S1E).^19^ These data highlight that our culture system generates a highly enriched population of a distinct cell state that may represent a key decision point on the trajectory to forming fully mature myelinating oligodendrocytes.

**Figure 1.**
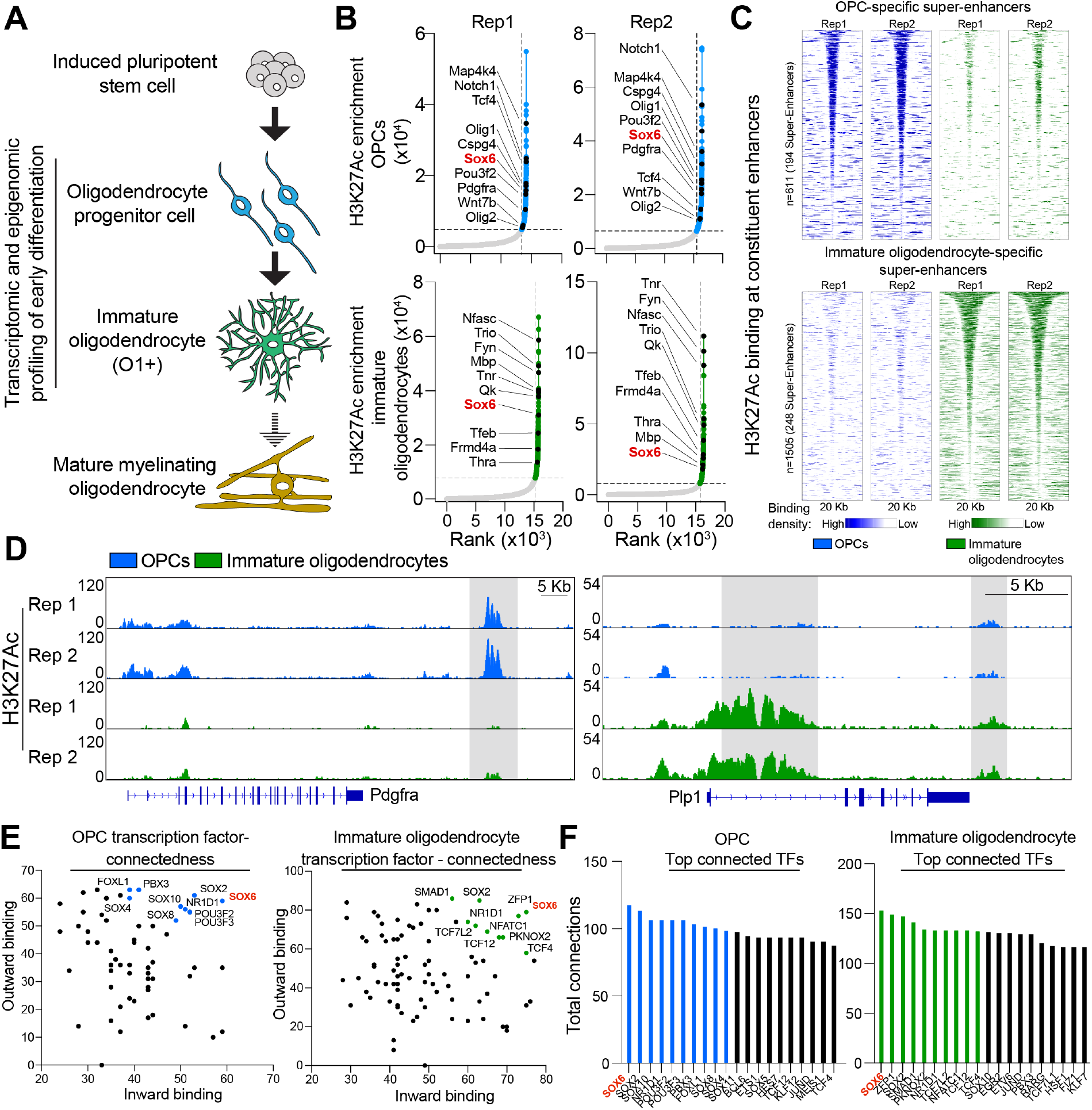
SOX6 is a Master Regulator of Immature Oligodendrocyte Formation. **(A)** Schematic depicting the formation of mature myelinating oligodendrocytes from mouse iPSCs. The stages profiled in the current study are indicated. **(B)** Hockey stick plots of input-normalized, rank-ordered H3K27Ac signal enrichment with super-enhancers highlighted in OPCs (in blue) and oligodendrocytes (in green). Example super-enhancer associated genes shared between replicates are listed and their associated regions are indicated as black circles. SOX6 is highlighted as a super enhancer associated transcriptional regulator in both replicates of both cell states. Data are presented as 2 biological replicates (two independent batches of OPCs from different mouse strains). See also Table S1. **(C)** Heatmaps of H3K27Ac signal in OPCs and oligodendrocytes at constituent enhancers of OPC-specific super enhancer loci (in blue) and at constituent enhancers of oligodendrocyte-specific super enhancer loci (in green). Data are presented as 2 biological replicates (two independent batches of OPCs from different mouse strains). **(D)** Genome browser view of two replicates of H3K27Ac ChIP-seq of OPCs (in blue) and oligodendrocytes (in green) at the locus for cell-type-specific super-enhancer-controlled genes *Pdgfra* and *Plp1*. Super-enhancer loci are highlighted in gray. Scale bars, 5Kb. **(E)** Scatter plot of the inward and outward binding for super-enhancer associated transcription factors in OPCs and immature oligodendrocytes. Each dot represents a single transcription factor and the top 10 connected transcription factors are indicated in blue (OPCs) or green (immature oligodendrocytes). SOX6 is highlighted in red as the most highly connected transcription factor. See also Figure S2 for total binding. **(F)** Bar graph of the top 50 transcription factors ranked by their total binding (sum of their inward and outward binding) in OPCs and immature oligodendrocytes. Each bar represents a single transcription factor and the top 10 transcription factors in terms of total binding are highlighted in blue (OPCs) and green (oligodendrocytes). SOX6 is highlighted in red as the top transcription factor for total binding in both states. See also Table S2. See also Figures S1 and S2

To find key regulators of fate decisions in OPCs and immature oligodendrocytes, we first identified super-enhancers, genomic regions defined by clusters of enhancers bound by master transcription factors (TF) known to drive expression of genes involved in cell identity (Figure 1B; Table S1).^26^ From H3K27Ac ChIP-seq datasets in both OPCs and immature oligodendrocytes, we identified super enhancers specific for OPCs or immature oligodendrocytes and their target genes (Figures 1C, 1D, S1F, S1G; Table S1). We also conducted motif analysis of accessible regions profiled by ATAC-seq to delineate cell-type-specific and shared TF motifs, such as *THRA* and *SOX* family motifs, respectively (Figure S1H; Table S2). Next, we integrated these data to perform transcriptional regulatory network analysis in OPCs and immature oligodendrocytes. We first identified super-enhancer regulated TFs. Then, for each TF identified, we computed TF connectedness by predicting the number of times it binds within enhancers of other superenhancer regulated TFs (outward binding) and the number of times other super-enhancer TFs bind to its own enhancers (inward binding) (Figure S2A; Table S3).^15,27,28^ We also calculated the prevalence of TF-binding motifs in regulatory networks assumed to be self-reinforcing based on integrating genome-wide active enhancers with regional data of focal DNA accessibility. Such networks are referred to as 'auto-regulatory cliques', and typically include master transcription factors (Figure S2A; Table S3).^15,27^ Here, our analyses identified SOX6 as the most connected transcription factor and central to auto-regulatory cliques, suggesting it may be a key regulator of early oligodendrocyte development (Figures 1E, 1F, S2B–S2D).

### miRNAs that accelerate oligodendrocyte development from OPCs converge on targeting SOX6

In parallel, we used genome-wide microRNA (miRNA) screening as an orthogonal approach to uncover transcriptional gatekeepers of oligodendrocyte development. In concert with transcription factors, miRNAs regulate the core circuitry governing cell states via post-transcriptional inhibition of many genes simultaneously, and therefore have substantial impacts on key transcription factor networks.^29,30^ Indeed, the miRNA processing machinery and multiple miRNAs have been shown to be critical for oligodendrocyte development.^18,31,32^ Thus, we conducted a genome-wide phenotypic miRNA screen consisting of 1309 miRNA mimics transiently transfected into OPCs, followed by high-content imaging and quantification of myelin basic protein (MBP), a protein enriched in maturing oligodendrocytes (Figure 2A, S3A, S3B and Table S4). In total, we identified 20 miRNAs that increased MBP expression in oligodendrocyte as single agents, including several miRNAs that are not endogenously expressed in the oligodendrocyte lineage (Figure 2A, 2B, and S3C). Top hits included miR-219 and miR-138, previously reported to be drivers of oligodendrocyte development in mice (Figure 2B and S3D).^31,32^ Through predictive targeting algorithms, we uncovered enriched gene targets of the miRNA hits, and identified *Sox6* as the top transcription factor target (predicted target of 8/20 miRNA hits), with no other highly connected transcription factor being significantly enriched (Figures 2C and S3E). Combined with our transcriptomic and epigenomic analyses, these functional data reveal that SOX6 is a key node in the circuitry governing oligodendrocyte development from OPCs.

**Figure 2.**
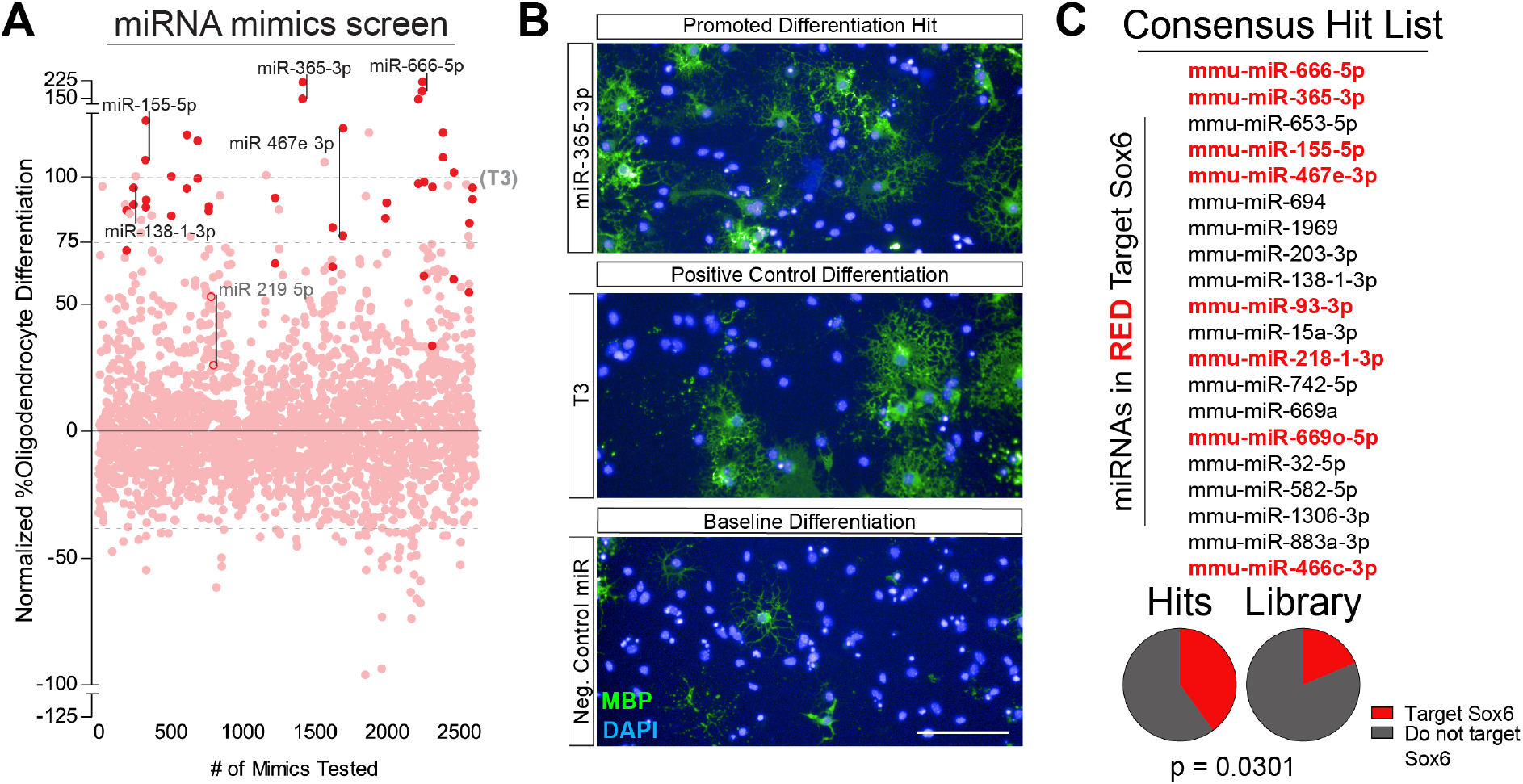
Exogenous miRNAs That Drive Oligodendrocyte Formation Converge on Regulating SOX6. **(A)** Primary miRNA mimic screen showing the effect of 1309 miRNAs on percentage of oligodendrocytes (MBP+ cells/ total DAPI) formed by OPCs relative to T3 treated OPCs. Each dot represents a single miRNA mimic in a single well of a 96 well plate and each miRNA mimic is present twice in biological duplicate with miRNA hits highlighted. Example miRNA hits and positive control miRNAs (mir-138 and miR-219) are labeled with a bar between replicates. See also Table S4 for the full list of miRNAs and their impact on oligodendrocyte formation. **(B)** Representative immunocytochemistry images of oligodendrocytes (MBP+ in green) from the primary screen of a top miRNA hit (miR-365-3p), T3 positive control, and negative miRNA mimic control. Nuclei are marked by DAPI (in blue). Scale bars, 100μm. **(C)** List of miRNA mimic hits that were called from the primary screen as miRNAs that drive oligodendrocyte formation from OPCs. miRNAs highlighted in red are those predicted to target SOX6. Pie charts indicate the number of miRNAs predicted to target SOX6 (in red) to miRNAs that do not target SOX6 (in dark gray) within top miRNA hits compared to their prevalence in the whole miRNA screening library.^56^ p-value was calculated using hypergeometric analysis. See also Figure S3.

### Loss of SOX6 accelerates oligodendrocyte maturation

Previous *in vivo* loss-of-function and *in vitro* studies in mice suggest that SOX6 maintains OPCs in the progenitor state by inhibiting the expression of oligodendrocyte genes and driving the expression of the OPC signature gene *Pdgfra*.^18,33,34^ Here, we found that knockdown of *Sox6* mRNA in OPCs led to a modest increase in the percentages of O1 and MBP positive oligodendrocytes at day 3 of differentiation (Figures 3A–3C, S4A, S4B). However, the morphology of the resulting oligodendrocytes was striking: a complex and matted myelin membrane indicative of enhanced maturation (Figures 3A–3C).^35^ Performing a time course experiment following SOX6 knockdown at days 1, 2, and 3 of differentiation, we observed that this precociously mature morphology was present by day 2 of differentiation, and was consistently more pronounced than the marginal increase in number of oligodendrocytes (Figures 3D–3F). Taken together, our observations indicate that SOX6, as opposed to regulating differentiation from the OPC state, may instead act as a gatekeeper of oligodendrocyte maturation.

**Figure 3.**
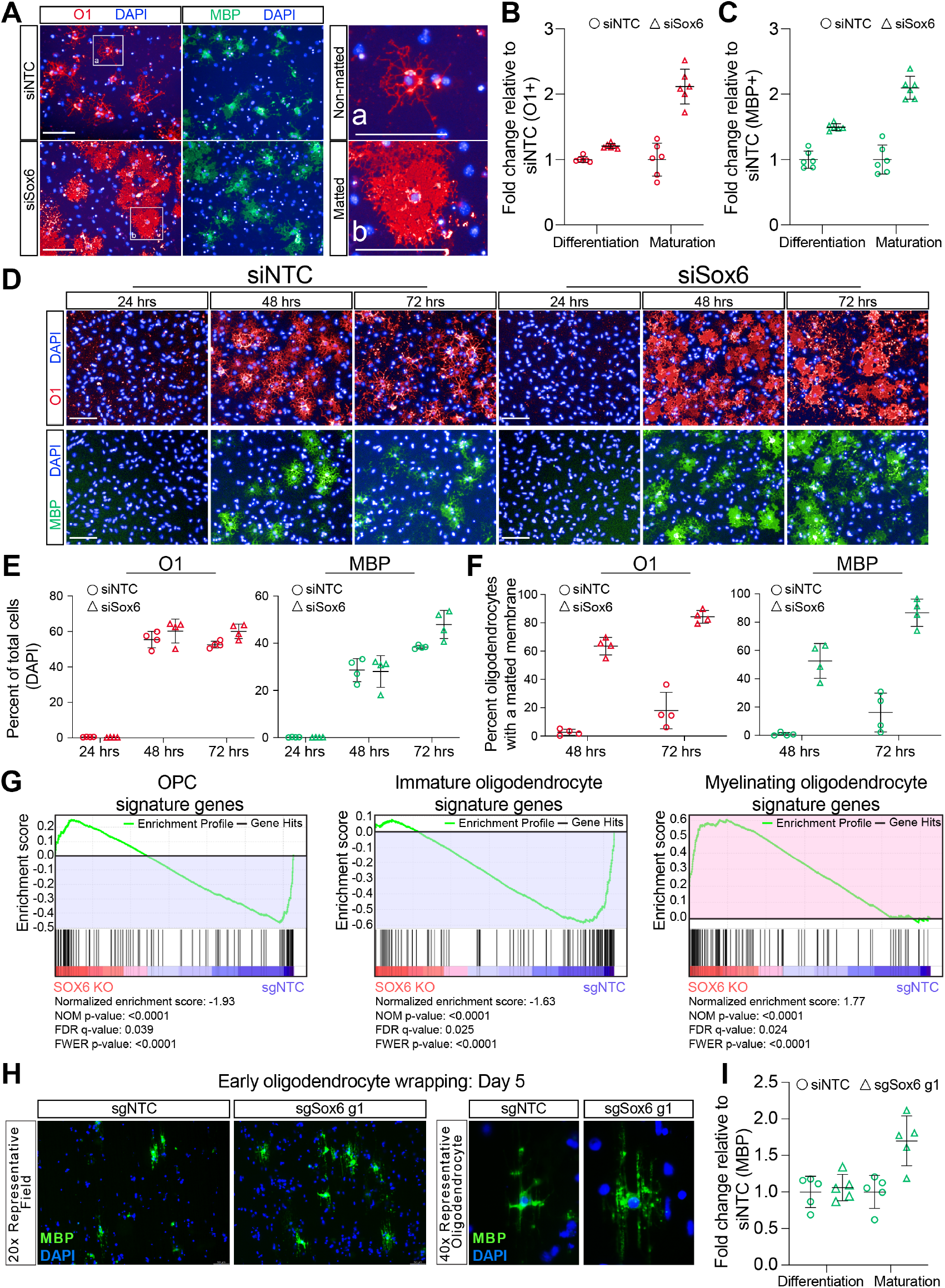
Loss of SOX6 Accelerates Oligodendrocyte Maturation. **(A)** Representative immunocytochemistry images of oligodendrocytes (MBP+ in green and O1+ in red) from OPCs transfected with non-targeting (siNTC) or SOX6 (siSox6) siRNAs. Nuclei are marked by DAPI (in blue). White boxes and zoomed in images to the right are examples of non-matted oligodendrocytes (a), and matted oligodendrocytes (b). Scale bars, 100μm. **(B-C)** Quantification of the percentage of oligodendrocytes of total cells (differentiation) and percentage of matted oligodendrocytes (maturation) of all oligodendrocytes for markers O1 (B) and MBP (C) from OPCs transfected with non-targeting (siNTC as circles) or SOX6 (siSox6 as triangles) siRNAs. Data are normalized to siNTC and presented as mean ± SD from 6 technical replicates (individual wells) from a single experiment. **(D)** Representative images of immature (O1+ in red) and late (MBP+ in green) oligodendrocytes at days 2 and 3 of differentiation from OPCs transfected with nontargeting or SOX6 siRNAs. Nuclei are marked by DAPI (in blue). Scale bars, 100μm. **(E)** Quantification of the percentage of O1 positive (in red) and MBP positive (in green) oligodendrocytes of total cells (indicated by total DAPI) from OPCs transfected with nontargeting (siNTC as circles) or Sox6 (siSox6 as triangles) siRNAs at days 1, 2, and 3 of differentiation. Data are presented as mean ± SD from 4 technical replicates (individual wells) from a single experiment. **(F)** Quantification of the percentage of oligodendrocytes displaying matted O1 (in red) and matted MBP (in green) from OPCs transfected with non-targeting (siNTC as circles) or SOX6 (siSox6 as triangles) siRNAs at day 2 and 3 of differentiation. Data are presented as mean ± SD from blinded quantification of images from 4 technical replicates (individual wells) from a single experiment. **(G)** Gene set enrichment analysis (GSEA) analysis of *in vivo* OPC (left), immature oligodendrocyte (middle), and myelinating oligodendrocyte (right) signature genes between SOX6 knockout (sgSox6 g1 and sgSox6 g2) and control (sgNTC) oligodendrocytes at day 3 of differentiation. This demonstrates a significant depletion of OPC and immature oligodendrocyte genes (highlighted in blue) and enrichment of myelinating oligodendrocyte genes (highlighted in red) in SOX6 knockout oligodendrocytes compared to control oligodendrocytes. See also Table S6 for the list of *in vivo* signature genes. **(H)** Representative immunocytochemistry images of early myelinating oligodendrocytes (MBP+ in green) on microfibers with all cells labeled with DAPI (in blue) at days 5 during differentiation from OPCs. Scale bars, 50 μm. **(I)** Quantification of the percentage of oligodendrocytes (Differentiation, MBP+ cells / total cells DAPI) and total MBP+ area of myelinating oligodendrocytes normalized to the total number of MBP+ cells (maturation) for sgNTC (circles) and sgSox6 g1 (triangles) at day 5 of differentiation on microfibers. Data are normalized to sgNTC and presented as mean ± SD from 5 separate microfiber wells per condition. See also Figure S4.

To investigate if the loss of SOX6 promotes transcriptional maturation of oligodendrocytes, we next generated SOX6 knockout OPCs. We used CRISPR-Cas9 with two independent single guide RNAs (sgSox6 g1 and sgSox6 g2), which led to a strong reduction of SOX6 protein expression compared to the non-targeting control (sgNTC) (Figure S4C). In contrast with expression profiles of non-targeting control *in vitro*-generated oligodendrocytes, RNA-seq of SOX6 knockout OPCs differentiated into oligodendrocytes revealed increased expression of the *in vivo* mature myelinating oligodendrocyte gene expression program, (Figures 3G and S4D–S4I; Table S5). These striking results suggest that loss of SOX6 unlocks cellular maturity and enables expression of transcriptional programs of mature myelinating oligodendrocytes.

Morphologic and transcriptional changes during maturation result in cellular adaptations that support mature function.^1^ To test how SOX6 suppression impacts mature oligodendrocyte functions, we used an established *in vitro* microfiber myelination assay.^36^ To this end, we transfected OPCs with miRNA mimics from our top screen hits (miR-666-5p and miR-365-3p) or siRNAs validated to target *Sox6* mRNA and assessed myelination at day 14. Each significantly increased the extent of microfibers myelinated as measured by high content imaging analysis of MBP immunostaining relative to controls (Figures S4J–S4M). Evaluation at an earlier timepoint (day 5) revealed that the loss of SOX6 had little effect on the percentage of oligodendrocytes formed but instead led to an increase in MBP+ myelin membrane per cell, which mirrored the 2-dimensional increase in membrane surface area seen with SOX6 suppression (Figures S4M, 3H, and 3I). Collectively, these data highlight a novel mechanism by which SOX6 governs the rate of oligodendrocyte maturation to form morphologically, transcriptionally, and functionally mature myelinating oligodendrocytes.

### SOX6 is enriched at super enhancers in OPCs and redistributes to cluster across gene bodies in immature oligodendrocytes

We next sought to understand the mechanism by which a lineage transcription factor such as SOX6 can perform seemingly separate functions across the post-mitotic oligodendrocyte lineage. To understand how SOX6 regulates oligodendrocyte maturation, we performed ChIP-seq of SOX6 in OPCs and immature oligodendrocytes. We found that SOX6 binds to nearly every superenhancer in OPCs and that the number of SOX6 binding peaks declines in immature oligodendrocytes in agreement with *Sox6* mRNA levels (Figures 4A, 4B, S5A and S5B). SOX6 was also strongly bound to its own locus in OPCs, which agrees with the self-regulating nature of master lineage transcription factors (Figure S5C and S5D).^15,27^ As OPCs differentiate, *Sox6* mRNA expression decreases, and we therefore expected SOX6 binding to simply decline or disappear in immature oligodendrocytes (Figure S5A and S5B). We were therefore surprised to see that SOX6 binding actually increased significantly at some loci, suggesting that SOX6 may redistribute in the immature oligodendrocyte state (Figures 4A and 4C). Many of the immature oligodendrocyte-specific SOX6 sites displayed a particularly interesting binding behavior in which the SOX6 signal spreads across gene bodies of genes that dramatically increase in expression in immature oligodendrocytes, such as *Bcas1* (Figures 4D and 4E). By ranking genes based on the average SOX6 intensity across their gene bodies, we identified and labeled the top regions of SOX6 pile-up as “SOX6 cluster genes,” which were similarly enriched for active and open chromatin across the entire gene body (Figure 4F, 4G, and S5E; Table S6). When we plotted the expression of these genes using multiple *in vivo* oligodendrocyte development datasets, we noticed that these genes are specifically induced in immature oligodendrocytes and then decline during maturation into myelinating oligodendrocytes (Figures 4H, 4I, and S6A–S6G; Table S5). In agreement, pathway analysis of SOX6-cluster genes demonstrated a clear enrichment for oligodendrocyte differentiation pathways (Figure 4J). Furthermore, loss of SOX6 significantly decreased SOX6-cluster gene expression in differentiating oligodendrocytes (Figure 4K). Collectively, these experiments suggest that SOX6 redistributes from extensive binding of superenhancers in OPCs to clustering across select gene bodies that are dramatically and transiently upregulated in immature oligodendrocytes.

**Figure 4.**
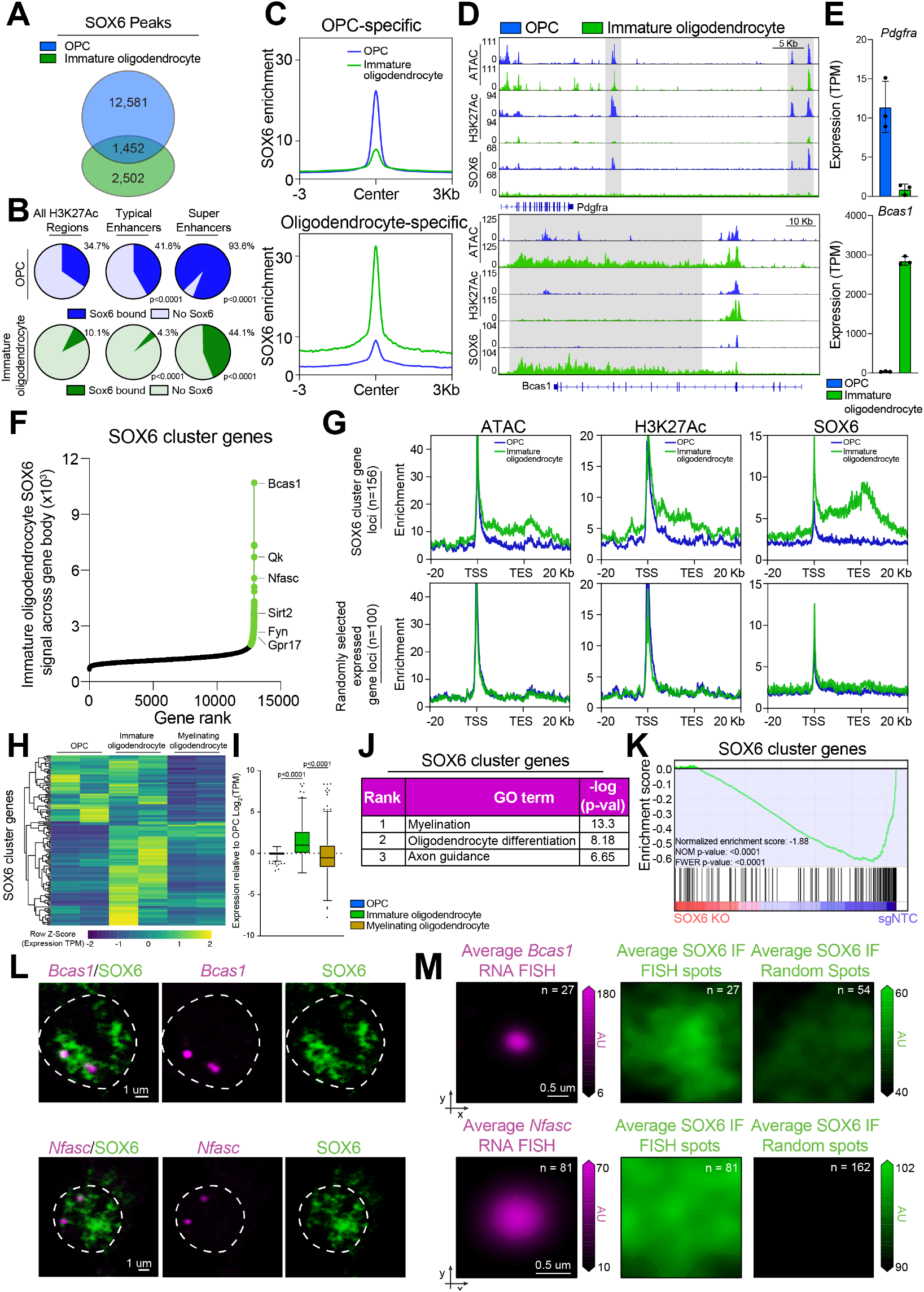
SOX6 Redistributes from OPC Super Enhancers to Cluster Across Gene Bodies in Immature Oligodendrocytes. **(A)** Venn diagram indicating the overlap between SOX6 peaks in OPCs (in blue) and immature oligodendrocytes (in green). **(B)** Pie charts indicate intersection of SOX6 peaks with all H3K27Ac peaks (FDR<0.001), typical enhancers, or super enhancers in OPCs (in blue) or oligodendrocytes (in green). p-values were calculated for typical enhancers or super enhancers compared to intersection with all H3K27Ac peaks using hypergeometric analysis. **(C)** Aggregate binding of OPC-specific SOX6 peaks (n= 12,581) and oligodendrocyte-specific SOX6 peaks (n=2,502) within 3Kb of the center of each peak in OPCs (in blue) and oligodendrocytes (in green) called by MACS2 (narrow peaks, FDR<0.001) normalized to input. **(D)** Genome browser view of ATAC-seq, H3K27Ac ChIP-seq, and SOX6 ChIP-seq in OPCs (in blue) and oligodendrocytes (in green) at SOX6 super-site associated loci including OPC-specific *Pdgfra* and Oligodendrocyte-specific *Bcas1*. SOX6 super-sites are highlighted in gray. Scale bars, 5Kb and 10Kb respectively. **(E)** Quantification of the normalized number of transcripts (TPM) for both *Pdgfra* and *Bcas1* in OPCs (in blue), and pre-myelinating oligodendrocytes (in green). Data represent mean ± SD from 3 biological replicates (independent samples) from RNA-seq. **(F)** Hockey stick plot of SOX6 signal across the gene body of all expressed genes in immature oligodendrocytes. Genes highlighted in green are those called as SOX6 cluster genes (SOX6 signal greater than 2-fold in immature oligodendrocytes compared to input and OPC). Example SOX6 cluster genes are listed. See also Table S5. **(G)** Aggregate plots of ATAC-seq, H3K27-Ac ChIP-seq, and SOX6 ChIP-seq across the gene bodies of SOX6 cluster genes (top) or 100 randomly selected expressed genes (bottom) and 20Kb upstream from the transcription start site (TSS) and 20Kb downstream from the transcription end site (TES). **(H)** Heatmap representation of row normalized expression of SOX6 cluster genes (TPM) in *in vivo* OPCs, immature oligodendrocytes, and myelinating oligodendrocytes. **(I)** Box and whisker plot of change in gene expression (TPM) relative to OPCs for SOX6 cluster genes in *in vivo* OPCs, immature oligodendrocytes, and myelinating oligodendrocytes. The black line represents the median with the box borders representing the upper and lower quartiles with dots representing statistical outliers. p-values were calculated using the Kruskal Wallis One-Way ANOVA with Dunn's multiple comparisons test. **(J)** Gene ontology (GO) analysis of genes associated with SOX6 clusters in immature oligodendrocytes. The chart includes curated pathways with their rank based on their respective p-values. See also Table S5 for full list of pathways. **(K)** Gene set enrichment analysis (GSEA) analysis of the SOX6 cluster genes in *in vitro* SOX6 KO compared to control immature oligodendrocytes demonstrates a significant depletion of SOX6 cluster genes in SOX6 KO immature oligodendrocytes. **(L)** Representative images of overlap between IF of SOX6 (green) and nascent RNA FISH of *Bcas1* and *Nfasc* (magenta) in fixed immature oligodendrocytes. **(M)** Average RNA FISH signal (magenta) and average SOX6 signal centered on RNA FISH foci with randomly assorted images of SOX6 staining used as a control. See also Figure S5.

### SOX6 clusters in immature oligodendrocytes represent gene melting events

Next, we sought to understand whether the SOX6 clusters could represent large transcriptional hubs that stabilize the post-mitotic intermediate oligodendrocyte state to ultimately control oligodendrocyte maturation over time. Specifically, the morphology of the chromatin tracks depicting SOX6 aggregation across gene bodies is mirrored by a recently described phenomenon known as “gene melting,” in which decondensed chromatin is observed across gene bodies of highly abundant, long genes.^37^ We therefore hypothesized that the SOX6 cluster loci represent gene melting events resulting from massive protein aggregation at genes being robustly expressed in immature oligodendrocytes. In agreement with this paradigm, genes harboring SOX6 aggregations in immature oligodendrocytes were significantly upregulated in our *in vitro* immature oligodendrocytes compared to OPCs (Figure S6G). In addition, such genes were significantly longer with an increased number of exons compared to all other expressed genes in immature oligodendrocytes (Figure S6H). SOX6 is an inherently disordered protein that appears to cluster across robustly expressed genes to promote gene melting and stabilize the immature oligodendrocyte state (Figure S6I).^37,38^ Performing high resolution confocal microscopy of immunofluorescence coupled with RNA-FISH confirmed SOX6 foci in immature oligodendrocytes that co-localized with the predicted SOX6-cluster genes *Bcas1* and *Nfasc* (Figure 4L and 4M). Of note, staining of these foci was more diffuse than staining of components of transcriptional phase condensates in other cell types, and RNA fish loci for *Bcas* and *Nfasc* demonstrated an elongated structure, which has been shown to characterize gene melting events (Figures 4L and S6J).^37,39^ Taken together, these data demonstrate that SOX6 forms transcriptional hubs at long, robustly expressed genes that are transiently expressed in immature oligodendrocytes.

### SOX6-regulated immature oligodendrocyte state is enriched in human neurological disease

Damage to myelinating oligodendrocytes in numerous neurological diseases leads to substantial disability, which is exacerbated by the impaired regenerative potential of residual OPCs that are unable to fully mature into myelinating oligodendrocytes.^22,40,41^ Using publicly available human single-nucleus RNA-seq datasets, we demonstrate that SOX6-cluster genes are enriched in cell populations found in multiple sclerosis patients compared to healthy controls, whereas there was no significant enrichment in Alzheimer’s disease(Figure 5A–5G).^42–44^ In contrast, transcripts from SOX6-cluster genes were depleted in Parkinson’s disease samples compared to controls (Figure 5A).^45^ These clinical observations suggest disease-specific stalling of the oligodendrocyte lineage in multiple sclerosis patients and highlight the potential for recovering white matter integrity with maturation-promoting therapeutics. Together, these data demonstrate that there is a disease-specific impact on different stages of the oligodendrocyte lineage and stresses the need for therapeutics that can promote the terminal progression of immature oligodendrocytes to functionally mature oligodendrocyte. Furthermore, our analyses highlight that maturation is a discrete and disease-relevant phase of development.

**Figure 5.**
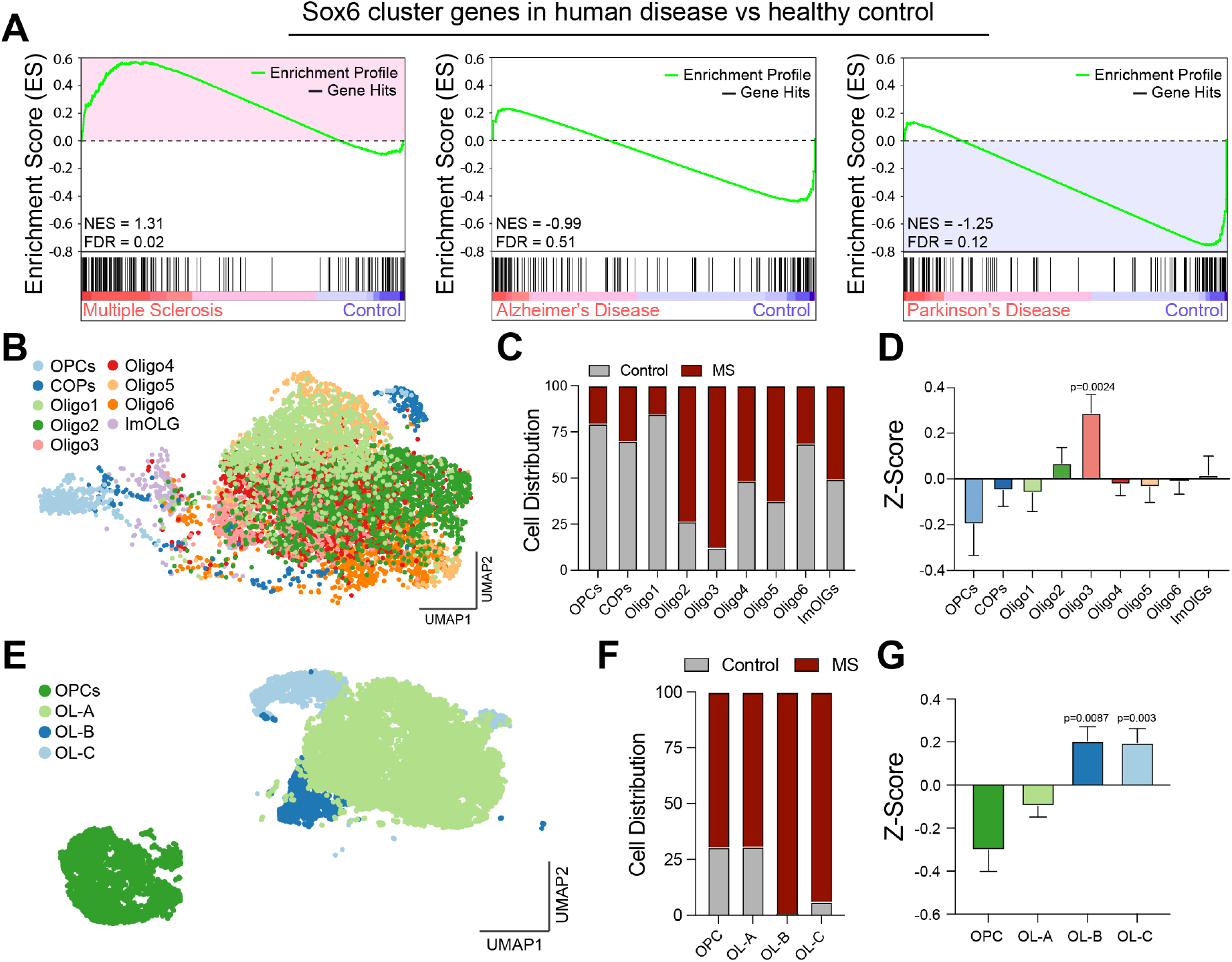
SOX6-regulated Immature Oligodendrocytes are Perturbed in Human Neurologic Disease. **(A)** Gene set enrichment analysis (GSEA) analysis of pre-ranked log2FC values of SOX6 aggregation genes between Multiple sclerosis, Alzheimer’s disease, and Parkinson’s disease and their respective healthy controls using publicly available data.^42–45^ This demonstrates a significant enrichment of SOX6 cluster genes in Multiple sclerosis (highlighted in blue) and depletion of SOX6 cluster genes in Parkinson’s disease (highlighted in red) compared to healthy control patients. **(B)** UMAP clustering of the oligodendrocyte lineage from multiple sclerosis and healthy control patients from publicly available data with individual clusters indicated.^42^ COP = committed oligodendrocyte progenitors, ImOLG = immune oligodendroglia, Oligo = oligodendrocyte. **(C)** Distribution of cells from multiple sclerosis and healthy control patients within each individual oligodendrocyte cluster from publicly available data.^42^ **(D)** Z-score normalized expression of SOX6 cluster genes in each of the oligodendrocyte clusters from publicly available data.^42^ Significant p-values are noted and were calculated using the Wilcoxon test. **(E)** tSNE clustering of oligodendrocytes from multiple sclerosis and healthy control patients from an additional publicly available dataset with individual clusters indicated.^43^ OL = oligodendrocyte cluster. **(F)** Distribution of cells from multiple sclerosis and healthy control patients within each individual oligodendrocyte cluster from publicly available data.^43^ **(G)** Z-score normalized expression of SOX6 cluster genes in each of the oligodendrocyte clusters from publicly available data.^43^ Significant p-values are noted and were calculated using the Wilcoxon test.

## DISCUSSION

During development, progenitor cells progress through differentiation and subsequent maturation to ultimately form functional tissues and organs.^1,16^ Although the fully differentiated cell state has generally been considered the final state transition for progenitor cells, it is becoming increasingly clear that the transition from a differentiated to a fully mature cell state is a highly regulated and distinct process.^1^ Accordingly, relative to the vast literature on directed differentiation approaches, the biological processes that control maturation are virtually unknown, and likely to be exquisitely customized in different organs and specialized cell types. There is a clear need to better understand maturation, as *in vitro* systems generally fail to move cells past immature states into fully functional, physiologically competent cells and tissues.^1–3,5–8,11^

In this study, we leveraged our tractable oligodendrocyte differentiation platform to gain insight into general biological processes that regulate cellular maturation.^25^ To accomplish this goal, we investigated transcription networks that drive both the formation and maintenance of the immature oligodendrocyte state. We focused our attention on transcription factors and microRNAs, as both can govern gene networks that control cell state. Through genome-wide interrogation of the mRNA and miRNA transcriptomes and the epigenomes of OPCs and immature oligodendrocytes, we successfully identified key regulatory nodes. By combining this rigorous profiling with a full genome functional miRNA screen, we were able to pinpoint SOX6 as a core transcription factor uniquely capable of broadly governing oligodendrocyte development.

Constitutive knockout of SOX6 in OPCs was previously reported to generate precocious myelin transcripts and proteins in mice; however, myelin formation in white matter tracts was dramatically impaired.^34^ This phenotype is likely the result of impaired migration of OPCs lacking SOX6 due to precocious maturation, which prevents them from reaching their proper locations within white matter tracts to form oligodendrocytes and mature in their correct destination.^34^ Functionally, SOX6 is thought to antagonize the pro-differentiation effect of SOX10 on oligodendrocyte formation such that loss of SOX6 enables precocious oligodendrocyte differentiation.^18,33^ However, we demonstrate that while loss of SOX6 has an effect on differentiation, the predominant effect was precocious formation of morphologically and transcriptionally mature oligodendrocytes. When we interrogated the whole-genome binding profile of SOX6, we found that SOX6 does not simply disappear during differentiation, but instead dramatically redistributes to form diffuse clusters at genes that are robustly and transiently expressed in intermediate, immature oligodendrocytes.

Upon close examination of the chromatin architecture using high resolution microscopy, we concluded that these loci may represent gene melting, a phenomenon describing chromatin decondensation at long, highly expressed genes that occurs in a cell-type-specific manner predominantly during development.^37,39,46^ Similar to previous studies, we demonstrate that these melting events represent robust open chromatin signatures across gene bodies.^37^ However, the potential transcriptional regulation of these melting loci and putative role in development remain unknown. Here, we demonstrate that dominant transcription factors, such as SOX6, can form transcriptional hubs underlying gene melting events that stabilize immature states within differentiation and maturation trajectories. This proposed mechanism explains how dominant lineage transcription factors are able to correctly instruct temporal regulation of maturation in post-mitotic cells. More broadly, identification of transcriptional regulators of gene melting in other cell lineages, such as pancreatic beta cells and red blood cells, two cell types of strong therapeutic interest that have been notoriously difficult to differentiate into fully mature states *in vitro*, could inform development of novel therapies capable of regenerating mature cells across various disease indications.^11,47^

OPCs proliferate and migrate to lesioned areas in multiple sclerosis patients but are unable to form mature myelinating oligodendrocytes.^22,24,40^ Here, we demonstrate that transcripts of SOX6-cluster gene targets are enriched in human multiple sclerosis brain tissue compared to healthy controls. These observations suggest that transient downregulation of SOX6 could offer a therapeutic approach in white matter disease that unlocks the terminal maturation of stalled intra-lesional oligodendrocytes. Mechanistically, our work dissects the transcriptional networks regulating oligodendrocyte formation and establishes a novel role for SOX6 as a governor of the rate of maturation by regulating gene melting. More broadly, our findings establish a novel mechanism by which transcriptional circuitry governs the rate of cell maturation along the continuum of cell states.

## ACKNOWLEDGEMENTS

This research was supported by grants from the National Institutes of Health R35NS116842 (P.J.T.), F30HD096784 (K.C.A), F30CA183510 (T.E.M.), F30CA236313 (A.R.M.), T32NS077888 (K.C.A., T.E.M., A.R.M., and M.S.E.), and T32GM007250 (K.C.A). M.A.S. was supported by a HHMI Hana Gray Fellowship and a NYSCF Druckenmiller Fellowship. Institutional support was provided by CWRU School of Medicine and philanthropic support was generously contributed by the Fakhouri, Long, Enrile, Peterson, Goodman, Geller, and Weidenthal families. Additional support was provided by the Small Molecule Drug Development, Genomics, and Light Microscopy and Imaging core facilities of the CWRU Comprehensive Cancer Center (P30CA043703) and the University of Chicago Genomics Facility. We are grateful to B. Zuchero and B. Barres for sharing materials, C. Lilliehook at Life Science Editors for editorial support, and D. Adams, Y. Federov, M. Madhavan, Y. Maeno-Hikichi, E. Shick, E. Cohn, D. Neu, W. Pontius, B. Gryder, and J. Kerschner for technical assistance and/or discussion.

## AUTHOR CONTRIBUTIONS

K.C.A., T.E.M., and P.J.T. conceived and managed the overall study. K.C.A., T.E.M., M.S.E., H.E.O., and L.R.H. performed, quantified, and analyzed *in vitro* experiments using mouse OPCs including qPCR, western blot, immunocytochemistry, miRNA screening, and generation of CRISPR knockout OPCs. K.C.A. performed ChIP-seq experiments with data analyses performed by A.R.M., K.C.A., T.E.M., D.C.F., and P.C.S. RNA-seq data analysis was performed by A.R.M., T.E.M., and K.C.A. J.K.V., T.E.M., and C.Y.L. performed transcriptional regulatory network analysis. K.C.A., M.A.S., J.E.H., and R.A.Y. designed, performed, and analyzed IHC+RNA FISH experiments. B.L.L.C. performed analyses of the immaturity signature in human diseases. K.C.A. assembled all figures. K.C.A. and P.J.T. wrote the manuscript with input from all authors.

## METHODS

### Culture of 293T Cells

293T cells (Takara Bioscience, 632180) were cultured in DMEM (Thermo Fisher, 11960-044) supplemented with 10% FBS (Fisher, A3160402), 1x MEM Non-essential amino acids (Thermo Fisher, 11140-050), 1x Glutamax (Thermo Fisher, 35050061), and 0.1 mM 2-Mercaptoethanol (Sigma, M3148). Media was changed every 48 hours.

### Pluripotent stem cell-derived OPC culture

Data were generated using OPCs generated from mouse epiblast stem cell (EpiSC) lines EpiSC5 (biological replicate 1 OPCs) and 129O1 (biological replicate 2 OPCs), which were derived from mouse strains 129SvEv and 129S1/SvImJ, respectively as described previously except SHH was not used for the maintenance or differentiation of OPCs.^25^ All OPC and immature oligodendrocyte cultures were maintained on plates or glass coverslips coated with 100 μg/mL poly(L-ornithine) (P3655, Sigma), followed by 10 μg/ml laminin (L2020, Sigma). OPC growth media consisted of DMEM/F12 supplemented with N2 Max (R&D Systems, AR009), B27 (Thermo Fisher, 12587010), 20ng/mL bFGF (R&D Systems, 23-3FB-01M), and 20ng/mL PDGFA (R&D Systems, 221-AA) and was changed every 48 hours. All cell and tissue cultures were maintained at 37° C with 5% CO2 in a humidified incubator. After 4 passages, these EpiSC derived OPCs were purified by fluorescence activated cell sorting using conjugated CD140a (eBioscience, 17-1401; 1:80) and NG2-AF488 (Millipore, AB5320A4; 1:100) antibodies. Sorted OPCs were then expanded and purity was verified by staining for markers of OPCs, astrocytes, and oligodendrocytes and frozen down in aliquots (Figure S1).

### Mouse OPC differentiation to oligodendrocytes

For immature oligodendrocyte formation, OPCs were seeded at 40,000 cells per well in 96-well plates or 50 million cells per 500 cm^2^ bioassay plate (166508, Thermo Fisher) (for generating large quantities of cells for sequencing studies) coated with PO and laminin. OPCs were plated in oligodendrocyte differentiation media consisting of DMEM/F12 supplemented with N2 Max, B27, 100ng/mL noggin (R&D, 3344NG050), 100ng/mL IGF-1 (R&D, 291G1200), 10uM cyclic AMP (Sigma, D0260-100MG), 10ng/mL NT3 (R&D, 267N3025), and 40ng/mL T3 (Thyroid hormone, Sigma, T-6397). Differentiation permissive media is identical to differentiation media except without the addition of T3. Differentiation proceeded for 3 days unless otherwise noted.

### Immunocytochemistry

For antigens requiring live staining (O1), antibodies were diluted in N2B27 media supplemented with 10% donkey serum (v/v) (017-000-121, Jackson ImmunoResearch) and added to cells for 18 minutes in a tissue culture incubator maintained at 5% CO_2_ and 37°C. Cells were then fixed with cold 4% paraformaldehyde (15710, Electron microscopy sciences) and incubated at room temperature for 18 minutes. Plates were washed three times with PBS and permeabilized and blocked in blocking solution, which consisted of 0.1% Triton X-100 in PBS supplemented with 10% normal donkey serum (v/v) (017-000-121, Jackson ImmunoResearch), for at least 30 minutes at room temperature. Primary antibodies were diluted in blocking solution and incubated on cells overnight at 4°C. Primary antibodies included anti-OLIG2 (1.2μg/mL, Proteintech, 12999-1-AP), anti-MBP (1:100, Abcam, ab7349), anti-O1 (1:50, CCF Hybridoma Core), anti-GFAP (1:5000, Dako, Z033401-2), anti-SOX6 (1:2000, abcam, 30455), anti-A2B5 (2μg/mL, Millipore, MAB312), anti-NKX2-2 (1:200, DSHB, 74.5A5), and anti-SOX10 (1:100, R&D, AF2864). The following day, cells were rinsed with PBS and incubated in blocking solution containing appropriate secondary antibodies conjugated to an Alexa-Fluor (4μg/mL, Thermo Fisher) and costained with DAPI (1μg/mL, D8417-1mg, Sigma).

### High content imaging and quantification

96-well plates and microfiber plates were imaged using the Operetta High Content imaging and analysis system (PerkinElmer). For 96-well plates, 8 fields were captured at 20x magnification per well unless noted otherwise. Images were then uploaded and analyzed with PerkinElmer Harmony and Columbus software as described previously.^21,23^ In brief, total cell number was identified using a threshold for area of DAPI staining of nuclei to exclude pyknotic nuclei and debris. To identify oligodendrocytes, each DAPI nucleus was expanded by 50% to determine potential intersection with staining of an oligodendrocyte marker (PLP1, O1, or MBP) in a separate channel. A threshold was set for each plate to determine whether expanded nuclei that intersected PLP1, O1, or MBP were scored as oligodendrocytes. The number of oligodendrocytes were then divided by the total number of cells as indicated by DAPI to give the percentage of oligodendrocytes per field. All fields were combined to give statistics on a per well basis.

### Generation of knockout OPCs

CRISPR knockout (KO) OPCs were created following a similar protocol used previously^23,48^. In brief, guides were selected from the Brie library^49^ including: sgNTC (AAGCCTACTTCACCGGTCGG), sgSox6 g1 (TTGACGGAATGAACTGTACG), and sgSox6 g2 (AGAACACGCTTTGAGAACCT). These were ordered as oligonucleotides from IDT, annealed, and cloned into the linearized CRISPRv2 backbone (Addgene, 52961).^50^ Clones were sequence verified by Sanger sequencing. 293T cells (Takara Bioscience, 632180) were then transfected using lenti-X shots following the manufacturer’s protocol (Clonetech, 631276). Transfection media was replaced with N2B27 base media after 24 hours. Lentivirus containing N2B27 was then collected after an additional 48 hours, filtered, supplemented with OPC growth factors PDGFA and FGF2, and added to OPCs at a ratio of 1:2 (v/v) with fresh OPC growth media. The next day, viral media was switched for fresh virus-free OPC growth media for 48 hours. Infected OPCs were then selected for 96 hours in OPC growth media supplemented with a lethal dose of puromycin (500ng/mL, Thermo Fisher, A1113802). OPCs were allowed to recover in selection free OPC growth media for at least 24 hours prior to being aliquoted and frozen down. For all experiments, infected CRISPR targeting and non-targeting control OPCs were derived from the same original batch of EpiSC-derived mouse OPCs. qPCR was performed to validate a reduction of gene targets for each batch of CRISPR KO OPCs generated.

### Magnetic sorting to purify oligodendrocytes

To purify oligodendrocytes for downstream applications such as RNA-seq, ChIP-seq, and ATAC-seq, we performed magnetic sorting using an antibody against O1 (Cleveland Clinic Hybridoma Core Facility)^19^, and followed the manufacturer’s protocol for the magnetic anti-mouse IgG secondary beads (Miltenyi, 130-048-402). In brief, OPCs were differentiated in oligodendrocyte differentiation media for 3 days and OPCs were cultured concurrently in OPC growth media. At the end of 3 days, oligodendrocytes and OPCs were live stained in suspension with anti-O1 (1:100, Cleveland Clinic Hybridoma Core Facility) with 10 million cells/mL in N2B27 supplemented with 1:15 BSA fraction V (Gibco, 15260-037) and 1:250 EDTA (Fisher, 324506-100ML) for 20 minutes on a rocker at 4°C. Cells were then washed with MACS solution consisting of 1:20 BSA (Miltenyi, 130-091-376) in autoMACs buffer (v/v) (Miltenyi, 130-091-222). Cells were then resuspended in MACS solution (80μl per 10 million cells) and magnetic anti-mouse IgG beads (20μl per 10 million cells) (Miltenyi, 130-048-402) and incubated on a rocker at 4°C for 20 minutes. Next, cells were washed, resuspended with MACS solution, strained to remove any clumps, and then processed by the autoMACS cell sorter using the “Posseld2” sorting option to obtain O1 positive and O1 negative populations. Oligodendrocytes were harvested from the O1 positive population from differentiation cultures whereas OPCs were harvested from the O1 negative population from OPC cultures. Enrichment for oligodendrocytes was apparent by RNA-seq and H3K27Ac ChIP-seq (Figures 1 and S1).

### Mouse oligodendrocyte microfiber myelination assay with siRNAs and miRNA transfection

Parallel-aligned 2-4 μm electrospun fibers fitted to 12-well plate inserts were placed in 12 well plates (AMSBio, AMS-TECL-006-4x). Prior to use, inserts were incubated in 70% ethanol. Next, fibers were coated with 100 μg/mL poly(L-ornithine) followed by 10 μg/ml laminin. Rep1 OPCs were seeded at 250,000 OPCs per well in differentiation permissive medium, allowed to attach for 2 hours, and were then transfected with 25 nM microRNA mimics from Horizon/Dharmacon (miR-365 mimic: C-310597-05-0002, miR-666 mimic: C-310691-01-0002), or siRNAs (Sox6: L-044291-01-0005), Non-Targeting Control: D-001206-14-05) using Dharmafect 3 siRNA transfection reagent (Dharmacon, T-2003-02). Media was replaced with normal differentiation permissive media 16 hours following the transfection. For T3 positive controls, medium was supplemented with 40ng/mL thyroid hormone from days 0-3. On day 3 thyroid hormone was removed, and medium was subsequently changed every third day. The cells were fixed at day 14 of differentiation by replacing media with 4% PFA for 15 minutes at room temperature followed by 3 PBS washes. All plates were permeabilized, blocked, and stained with primary antibody as described in the immunocytochemistry section. Primary antibodies included rat anti-MBP (1:100, Abcam, ab7349) and rabbit anti-Olig2 (1.2μg/mL, Proteintech, 12999-1-AP). The next day, cells were rinsed with PBS and incubated with the appropriate secondary antibody conjugated to an Alexa-Fluor (4μg/mL, Thermo Fisher) along with the nuclear stain DAPI (1μg/mL, D8417-1mg, Sigma). Plates were imaged on the Operetta^®^ High Content Imaging and Analysis system. A total of 30 fields were captured at 20x using Acapella^®^ software, and images were analyzed using Harmony^®^ software and Columbus™ software. Using this analysis software, we developed an Acapella^®^ script to quantify the total area of MBP+ oligodendrocytes normalized to the number of OLIG2 positive cells per well.

### Mouse oligodendrocyte microfiber myelination assay with sgNTC and sgSox6 g1 OPCs

96-well plates with parallel-aligned 2-4 μm electrospun fibers (AMS.TECL-005-8X, AMSBio) were washed with 70% ethanol followed by coating with PO for at least 1 hour at 37°C and laminin for at least 3 hours at 37°C. 40,000 sgNTC or sgSox6 g1 OPCs were then added per well in differentiation media taking care to avoid damaging the fibers towards the bottom of the well. Media was changed with fresh differentiation media plus penicillin-streptomycin (15070-063, Thermo Fisher) every 48 hours. After 5 days, cells were fixed, permeabilized, blocked, and stained with anti-MBP (1:100, Abcam, ab7349) and DAPI following the protocol outlined for immunocytochemistry. Plates were imaged on the Operetta^®^ High Content Imaging and Analysis system. A total of 8 fields were captured at 20x using Acapella^®^ software, and images were analyzed using Harmony^®^ and Columbus™ software. The total area of MBP+ oligodendrocytes per well was calculated using the same script for the 12-well microfiber inserts and the number of MBP+ oligodendrocytes were manually counted in each field and added together for per-well statistics. Differentiation was calculated by dividing the number of MBP+ oligodendrocytes by the total number of DAPI positive cells. Maturation was determined by quantifying the total area of MBP normalized to the total number of MBP+ oligodendrocytes.

### ChIP-seq and alignment to genome

OPCs and purified O1+ oligodendrocytes were harvested using the autoMACS automated cell sorter as described in the Magnetic sorting to purify immature oligodendrocytes (O1+) section. Fixation, nuclei isolation, and chromatin shearing were performed as previously described using the Covaris TruChIP protocol following the manufacturer’s instructions for the “high-cell” format.^23^ In brief, sorted OPCs and O1+ oligodendrocytes were crosslinked in “Fixing buffer A” supplemented with 1% fresh formaldehyde for 10 minutes at room temperature with oscillation and quenched for 5 minutes with “Quench buffer E.” Next, cells were washed with PBS and immediately proceeded to nuclei extraction. Isolated nuclei were then sonicated using the Covaris S2 with 5% Duty factor, 4 intensity and four 60-second cycles. Sheared chromatin was then cleared and incubated with protein G magnetic DynaBeads (Thermo Fisher, 10004D) that had been pre-incubated with primary ChIP-grade antibodies overnight at 4°C. Primary antibodies used included anti-H3K27Ac (9μg/sample, Abcam, ab4729), and anti-SOX6 (15 μg/sample, Abcam, ab30455). Protein DynaBeads were then washed, and DNA was eluted, reverse cross-linked, and treated with RNAse A followed by Proteinase K digestion. ChIP DNA was purified by phenolchloroform separation and used to construct Illumina sequencing libraries that were sequenced on the HiSeq2500 or NextSeq with single-end 50bp or 75bp reads respectively with at least 20 million reads per sample.

For aligning reads to the genome, reads were quality and adapter trimmed using Trim Galore! Version 0.3.1. Trimmed reads were aligned to the mouse genome (mm10) with Bowtie2 version 2.3.2 and duplicate reads were removed using Picard MarkDuplicates. Peaks were called with MACSv2.1.1 with an FDR<0.001 using the broad peaks subcommand for histone marks (H3K27Ac) and narrow peaks subcommand for transcription factors (SOX6) and normalized to background input genomic DNA. Peaks were visualized using the Interactive Genomics Viewer (IGV, Broad Institute). Cluster plots were generated using deepTools2 computeMatrix followed by plotHeatmap (https://deeptools.readthedocs.io/en/develop/).^51^ Peaks were assigned to the nearest expressed gene (TPM>1) using bedtools closest function.^52^

### Super enhancer analyses

Super enhancers (SE) for OPCs and immature oligodendrocytes were called using the ROSE algorithm on H3K27Ac ChIP-Seq data. Super enhancers were called separately for biological replicates rep1 and rep2. Gene and miRNA targets of these SEs were called by linking the superenhancer to the closest expressed gene. Cell-type-specific super-enhancers were called using the dynamic enhancer algorithm (https://github.com/BradnerLab/pipeline/blob/master/dynamicEnhancer.py). State-specific superenhancers were then linked to the closest expressed gene targets that exhibited cell-type-specific expression (fold change > 2.5 in one state compared to the other).

### Omni ATAC-seq

Omni ATAC-Seq was performed on 50,000 OPCs and sorted O1+ oligodendrocytes based on the protocol described previously.^53^ In brief, nuclei were extracted from cells and treated with transposition mixture containing Nextera Tn5 Transposase (Illumina, FC-121-1030) for 30 minutes at 37°C with 1000 RPM mixing. Transposed fragments were then purified using Qiagen MinElute columns (Qiagen, 28004), PCR amplified, and libraries were purified with Agencourt AMPure XP magnetic beads (Beckman Coulter) with a sample to bead ratio of 1:1.2. Samples were sequenced on the HiSeq2500 with single-end 50bp reads with 100 million reads per sample. Reads were aligned to the mm10 mouse genome following the same pipeline that was used for aligning ChIP-seq data and peaks were called using MACS2 narrowpeak subcommand.

### Motif enrichment analysis

Motifs were called under significant SOX6 peaks (FDR<0.001) or ATAC-Seq peaks within super enhancers (FDR<0.001) using HOMERv4.11.1.^14^ The FindMotifsGenome.pl tool was used with 200bp windows for ATAC-Seq regions and SOX6 peaks using mm10 as the reference genome.

Motifs enriched under super-enhancers were called by intersecting super-enhancers between biological replicates, intersecting these super-enhancers with ATAC-seq peaks, and performing HOMER motif analysis under these regions as specified above. Motifs were considered significantly enriched if below a p-value cutoff of 1×10^-10^ and expressed in OPCs and/or immature oligodendrocytes. Significantly enriched motifs in OPCs and immature oligodendrocytes were then overlapped to provide the Venn diagram in Figure S1H.

### Calling SOX6-cluster genes

SOX6 intensity in OPCs and immature oligodendrocytes were tabulated across expressed gene bodies (from transcription start site to end site) in immature oligodendrocytes using the deepTool multiBigwigSummary. SOX6-cluster genes were called if the average intensity across the gene’s body was 1) greater than 2-fold over background (input) and 2) greater than 2-fold over SOX6 average intensity at these loci in OPCs. This resulted in 156 cluster gene loci. See also Table S5 for the list of SOX6 cluster genes.

### Bulk RNA-seq sample preparation and alignment

OPCs and immature oligodendrocyte cultures were lysed in TRIzol and RNA was isolated as described for qPCR. Specifically, RNA for defining in vitro OPC and immature oligodendrocyte expression profiles were generated from magnetically sorted OPCs and O1+ oligodendrocytes. For RNA isolated from sgNTC, sgSox6 g1, and sgSox6 g2 OPCs and immature oligodendrocytes, cells were plated into 6-well plates in OPC growth media and differentiation permissive media respectively for 60 hours at which point cells were lysed with TRIzol. Libraries were prepared following protocols from NEBNext Poly(A) mRNA Magnetic Isolation Module (NEB, E7490L) and NEBNext Ultra RNA Library Prep Kit for Illumina (NEB, E7530L). In brief, samples were enriched for mRNA using oligo(dT) beads, which were fragmented randomly and used for cDNA generation and subsequent second-strand synthesis using a custom second-strand synthesis buffer (Illumina), dNTPs, RNase H and DNA polymerase I. cDNA libraries then went through terminal repair, A-base ligation, adapter ligation, size selection, and PCR enrichment. Final libraries were pooled evenly and sequenced on the Illumina NovaSeq with paired-end 150bp reads with a read-depth of at least 20 million reads per sample.

For gene expression analysis, reads were aligned to the mm 10 genome and quantified in transcripts per million (TPM) values using salmon 0.14.1 (https://github.com/COMBINE-lab/salmon). Transcripts were summarized as gene-level TPM abundances with tximport. A gene with TPM>1 was considered expressed. Differential expression analysis was performed using DESEQ2 (https://bioconductor.org/packages/release/bioc/html/DESeq2.html). Significant genes were called based on p-adj and fold change values as described in the results section.

### miRNA screening

#### Plate preparation

96-well CellCarrier plates treated with poly-D-lysine (PerkinElmer) were coated with laminin (Sigma, L2020; 10 mg/ml) using electronic multichannel pipettors. Laminin was added to plate in 50 ul of base media (DMEM/F12 supplemented with N2 (R&D Systems), B-27 (Life Technologies) and incubated at 37°C for 30 minutes prior to plating cells.

#### Cell plating

Cells were added directly to wells without aspirating the laminin. 25,000 OPCs were seeded per well in 100 ul of 1.5x concentrated differentiation permissive media, creating a final volume of 150 uL and concentration of 1x differentiation permissive media (described above). Cells were allowed to attach for 2 h before transfection.

#### miRNA mimics/inhibitors and plate set-up

We used the mouse Dharmacon miRIDIAN miRNA Mimic/Inhibitor Bundle v19.0 (Thermo Fisher Scientific Biosciences). This bundle contained 1309 miRNA mimic and 1309 miRNA inhibitors towards all known mouse miRNAs as described by the miRbase v19.0 release, arrayed in 96 well plates. 80 miRNA mimics were arrayed across each 96 well plate from columns 2-11, leaving columns 1 and 12 open for negative and positive controls. Matched miRIDIAN miRNA negative controls were used in all 8 wells of column 1 in each plate as negative controls. Thyroid hormone was added at final concentration of 40ng/mL to cells in column 12 as a positive control.

#### miRNA mimic preparation

miRNA mimic plates contain 0.1 nm of lyophilized nucleic acid. These plates were spun at 4000 RPM for 2 min to pellet nucleic acid. These were diluted to 2 uM with by adding 50 uL of 1x PBS to each well in sterile conditions. Plates were then rocked for 90 minutes at RT to resuspend miRNA mimics.

#### Transfection

Each miRNA screening plate was screened in duplicate. Transfections were prepared for 2 plates using a separate 96 well plate. 2.75 uL of miRNA mimic dilution above was added to 55 uL 1x differentiation permission media in each well. 0.25 uL of Dharmafect #3 (Thermo Fisher) was used per well. 220 uL Dharmafect #3 was added to 22 mL of differentiation permissive media as a master mix. 52 uL of this mix was added to each well and mixed well before incubating at RT for 20 minutes. This is enough for transfections across two plates. Then, 100 uL of media was taken off of plated cells from above using a multichannel repeat pipettor, leaving 50 uL. 50 uL of transfection mixture for each well was added to plated cells to give a final volume of 100 uL in each well. Cells were incubated under standard conditions (37°C, 5% CO_2_) overnight. The next morning, 50 uL of media was removed and replaced with 100 uL of differentiation permissive media. Cells we then incubated for an additional 2 days.

#### Read out

3 days after transfection, cells were fixed with 4% paraformaldehyde (PFA) in phosphate buffered saline (PBS). Fixed plates were permeabilized with 0.1% Triton X-100 and blocked with 10% donkey serum (v/v) in PBS for 20 min. Cells were labelled with anti-MBP (1:100, Abcam, ab7349) in 10% donkey serum (v/v) in PBS for 1.5 h at room temperature (22°C) followed by detection with Alexa Fluor conjugated secondary antibodies (1:500) for 45 min. DAPI was added during washing steps to enable nuclei visualization (Sigma; 1 mg/ml). Plates were imaged and quantified using high content imaging described above.

### Small RNA-seq preparation and analysis

OPCs and immature oligodendrocyte cultures or primary cell isolations were lysed and processed using the miRNeasy Mini kit (Qiagen, 217004) according to manufacturer’s instructions. Primary cell isolations were generously provided by Dr. Ben Barres and Dr. Brad Zuchero. Libraries were prepared using the TruSeq Small RNA Sample Prep Kit (Illumina, RS-200-0112) following protocols from the preparation guide (RS-930-1012, 2011 Rev. C). In brief, samples were enriched for miRNA by utilizing the 3’ hydroxyl group and 5’ phosphate group found on most miRNAs to add a 3’ and 5’ adapters specific to these modifications. This was following by RT-PCR for 1^st^-strand synthesis and then PCR-amplification to add on library indexes. Finally, libraries were size selected by gel purification. Final libraries were pooled evenly and sequenced on the Illumina HiSeq 2500 with single-end 50bp reads.

The standalone version of sRNAbench version 05/2014 (https://bioinfo2.ugr.es/srnatoolbox/standalone/) was used to align reads to miRBase 20 miRNAs and the mm9 genome, allowing for prediction of novel miRNAs. The multi-mapping read alignment strategy was used. Expression of mature sense miRNAs was assessed, and levels were RPM normalized using the reads that mapped to the library as the baseline. Differential expression analysis was performed using DESEQ2 (https://bioconductor.org/packages/release/bioc/html/DESeq2.html). Significant genes were called based on p-adj and fold change values as described in the results section.

### Generation of *in vivo* signature gene sets

*In vivo* signature gene sets were generated using publicly available RNA-seq data (GEO: GSE52564) of *in vivo* purified OPCs, immature oligodendrocytes, and myelinating oligodendrocytes.^19^ Genes were first filtered for those expressed in our *in vitro* OPCs and/or immature oligodendrocytes (average TPM > 1). Next, the average expression (TPM) for each individual gene in a single cell type was divided by the average expression of the same gene in the other two cell types. These values were then ranked and the top 100 genes were selected as *in vivo* signature genes for the specified cell-type.

### Box and whisker plots

For box and whisker plots, RNAseq replicates were divided by the average of the respective control (TPM) on a per gene basis in each category and then all individual replicates were plotted together. Box and whisker plots were generated using the Tukey method in Prism GraphPad software to plot the whiskers and outliers.

### Gene ontology analysis

Metascape (http://metascape.org/) was used to identify significant pathways from desired gene lists. The pathway name, rank, and p-value are recorded in provided tables in the results.^54^

### Gene set enrichment analysis

Gene set enrichment analysis (GSEA) was performed for SOX6-cluster genes and *in vivo* signature gene sets using classic scoring, 1000 gene-set permutations, phenotype permutation, and signal-to-noise metrics. Normalized enrichment scores, NOM p-values, false discovery rate, and FWER p-values were calculated by GSEA software (https://www.gsea-msigdb.org/gsea/index.jsp).^55^

To analyze enrichment of SOX6-cluster genes in disease, pre-ranked Log2FC values were generated for GSEA analysis using Seurat to subset oligodendrocyte lineage cells from publicly available single-nuclei data sets from multiple sclerosis^42,43^, Alzheimer’s^44^, and Parkinson’s^45^ patients (GEO: GSE118257, PRJNA544731, GSE138852, and GSE157783 respectively). Then the Log2 fold-change between patient and healthy control cells was calculated using the Seurat command *FoldChange(*). The Log2FC values were then ranked from greatest to smallest and used as input for GSEA. GSEA pre-ranked analysis was ran using GSEA v4.1 software from the Broad Institute with default settings.

### Human single-nucleus RNA-seq analysis and data visualization

UMAP and tSNE plots were generated using published embeddings for single nuclei RNAseq data (GEO: GSE118257 and PRJNA544731) from UCSC Cell Browser and colored by published oligodendrocyte lineage cell subtypes using Seurat. The distribution of nuclei from multiple sclerosis patients or healthy controls was calculated within each of the oligodendrocyte lineage clusters.^42,43^ Seurat was used to calculate the average expression of SOX6-cluster genes within each oligodendrocyte lineage cluster and then a Z-score was generated for each gene across the oligodendrocyte lineage subtypes.

### Predicting targets of miRNAs

Predicted target genes for a given miRNA and vice versa were called using mirPath v.3 using the microT-CDS option with a microT threshold of 0.65 and p-value threshold of 0.05.^56^ This was used to tabulate the number of super-enhancer miRNAs that were predicted to target individual members of the transcriptional regulatory network in OPCs and immature oligodendrocytes. This was also used to determine whether a transcription factor in the OPC and immature oligodendrocyte transcriptional regulatory network was significantly enriched as a putative target of miRNA mimic hits from the phenotypic miRNA screen. In brief, the fraction of miRNA mimic hits targeting the transcription factor (out of the 20 hits) was compared with the fraction of miRNA mimics in the whole screening library that target the transcription factor using hypergeometric analysis. p-values were reported for each transcription factor tested.

### Transcriptional regulatory network analysis

Transcriptional regulatory analysis was performed using H3K27Ac ChIP-seq, RNA-seq, and ATAC-seq data derived from sorted rep 1 OPCs and immature oligodendrocytes. In brief, COLTRON (https://pypi.org/project/coltron/) calculated inward and outward binding of superenhancer regulated transcription factors in both states. The presence of transcription factors in autoregulatory cliques (clique fraction) was also calculated.^15,27^

### Western blot

At least 1 million OPCs or O1+ sorted immature oligodendrocytes were collected and lysed in RIPA buffer (Sigma, R0278) supplemented with protease and phosphatase inhibitor (78441, Thermo Fisher) for at least 15 minutes and cleared by centrifugation at 13,000g at 4°C. Protein concentrations were determined using the Bradford assay (Bio-Rad Laboratories). Protein was diluted, boiled at 95°C for 5 minutes, run using NuPAGE Bis-Tris gels (NP0335BOX, Thermo Fisher), and then transferred to PVDF membranes (LC2002, Thermo Fisher). Membranes were blocked in 5% nonfat milk (Nestle carnation) in TBS plus 0.1% Tween 20 (TBST) for 30 minutes and then incubated with primary antibodies at 4°C overnight in blocking solution. Primary antibodies used included anti-SOX6 (1μg/mL, Abcam, ab30455) and B-Actin peroxidase preconjugated antibody (1:50,000, Sigma, A3854). Membranes were then imaged and analyzed using Licor Image Studio™ software. Westerns were normalized to the beta-actin loading control.

### qRT-PCR

At least 500,000 OPCs or oligodendrocytes were lysed in TRIzol (Ambion) followed by purification and elution of RNA using phenol-chloroform extraction and the RNeasy Mini Kit (74104, Qiagen). RNA quality and quantity were determined using a NanoDrop spectrophotometer. cDNA was generated using the iSCRIPT kit following the manufacturer’s instructions (1708891, Biorad). qRT-PCR was performed using pre-designed TaqMan gene expression assays (Thermo Fisher) including: *Sox6* (Mm00488393_m1) and *Rpl13a* (Mm05910660_g1). qPCR was performed using the Applied Biosystems 7300 real-time PCR system and probes were normalized to *Rpl13a* endogenous control.

### siRNA knockdown of *Sox6*

siRNAs used for knockdown of *Sox6* were purchased from Horizon/Dharmacon (Sox6: L-044291-01-0005, Non-Targeting Control: D-001206-14-05 5 nmol) and nucleofected into OPCs using the Basic Nucleofector Kit for Primary Mammalian Glial Cells (Lonza, VPI-1006) following manufacturer’s instructions. OPCs (5 million cells per nucleofection) were resuspended in nucleofection solution with 0.6μl of the desired 50μM siRNA solution, nucleofected using the Amaxa Nucleofector 2b (Lonza, AAB-1001) using the A033 setting, and then plated in differentiation media supplemented with penicillin-streptomycin (Thermo Fisher, 15070063) for the time indicated in the results. Remaining OPCs were plated in OPC growth media supplemented with penicillin-streptomycin (Thermo Fisher, 15070063) for 48 hours and then lysed with RIPA buffer (Sigma, R0278-500ML) supplemented with protease inhibitors (Thermo Fisher, 87786) and processed for western blot.

### Calculating oligodendrocyte maturation in 2D culture

OPCs treated with siNTC or siSox6 were fixed and stained for O1 and MBP and imaged using the operetta as described in the immunocytochemistry and high content imaging and quantification sections. To quantify observed matted vs non-matted oligodendrocyte morphology in an unbiased manner, 6 images per well (for Figures 3A–3C) and 4 images per well (for Figures 3D–3F) were randomly selected, and de-identified. A reviewer blinded to both the image identities as well as the hypothesis of the experiment was provided with examples of matted or non-matted oligodendrocytes and asked to quantify the number of matted and non-matted oligodendrocytes in each image. After cell counts were received, the results were re-aligned with their proper ID and the percentage of matted cells was calculated for each image and then averaged and reported on a per well basis.

### SOX6 disorder calculations

Disorder values were calculated for SOX6 (UniProt ID: P40645) using publicly available algorithms: VSL2 (http://www.pondr.com/) and IUPred2 and ANCHOR2 (https://iupred2a.elte.hu/).^57,58^

### Co-immunofluorescence with RNA FISH

OPCs were grown and differentiated on 24-well coverslips that had been polyornithine and laminin coated previously. Immunofluorescence was performed as described earlier using anti-SOX6 (1:1000) and anti-O1 (1:100) for primary antibodies. After secondary antibodies were washed off 3 times with PBS, cells were fixed again with 4% PFA in PBS for 10 minutes at room temperature. RNA fish was then performed per Stellaris protocol. In brief, 20% RNase-free Stellaris wash buffer A (Biosearch Technologies, SMF-WA1-60), 10% deionized formamide (EMD Millipore, SS4117), and 70% RNase-free water was added for 5 minutes at room temperature. Coverslips were that moved to hybridization buffer containing 90% Stellaris hybridization buffer (Biosearch Technologies, SMF-HB1-10) and 10% deionized formamide, and 12.5uM Stellaris custom nascent Quasar 570 RNA FISH probes (targeting intronic regions of top SOX6 cluster genes *Bcas1* or *Nfasc*) (Biosearch Technologies, SMF-1063-5) overnight in the dark at 37°C. The following day, the coverslips were washed and nuclei were stained with DAPI followed by mounting onto slides using prolong glass antifade mountant (Thermo Fisher, P36984). Images were acquired using an RPI Spinning Disk confocal microscope with a 100x objective. RNA FISH probes were designed to target introns of *Bcas1* and *Nfasc* using Biosearch Technologies custom probe design software (www.biosearchtech.com/stellaris-designer) with masking level set to 5 and probe size of 20.

### Image analysis for co-immunofluorescence with RNA FISH

Multichannel z-stack images for coIF/RNA-FISH were analyzed using methods described in ^59^ First, images were maximally projected in the z plane. Nuclei were segmented using DAPI signal with a median filter (20 px) followed by automated Li thresholding using the Python library scikit-image. Nascent RNA-FISH spots were segmented using the RNA-FISH channel image. These images were Gaussian filtered (sigma=2), and spots were segmented by an automated threshold (3 standard deviations above the mean intensity of the image) and required to be in the nucleus. As a control, for each FISH spot detected, two spots in the nucleus were chosen by random pixel selection. For each FISH spot or random spot, a box (70×70 pixels) was extracted from the multichannel image, and intensities for both the FISH and IF signal were averaged across all detected FISH spots to generate contour plots.

### Statistics and replicates

GraphPad Prism was used to perform statistical analyses unless otherwise noted. Statistical tests and replicate descriptions are detailed in each figure legend. Black filled-in circles for bar graphs indicate biological replicates (independent experiments) whereas open circles represent technical replicates. Statistics were only performed on samples with biological replicates. Data was graphed as mean ± standard deviation (SD) or ± standard error of the mean (SEM) as detailed in the figure legend. A p-value less than 0.05 was considered significant unless otherwise noted.

### Data availability

Further information and requests for resources and reagents should be directed to and will be fulfilled by the lead contact, Paul Tesar. All datasets generated in this study have been deposited in Gene Expression Omnibus (https://www.ncbi.nlm.nih.gov/geo/) under SuperSeries accession code GSE197319 with subseries for RNA-seq (GSE181952), ChIP-seq (GSE182245), miRNA-seq (GSE183160), and ATAC-seq (GSE182558).

Access key: mjcxeaegnlahxop

---

**Figure S1.**
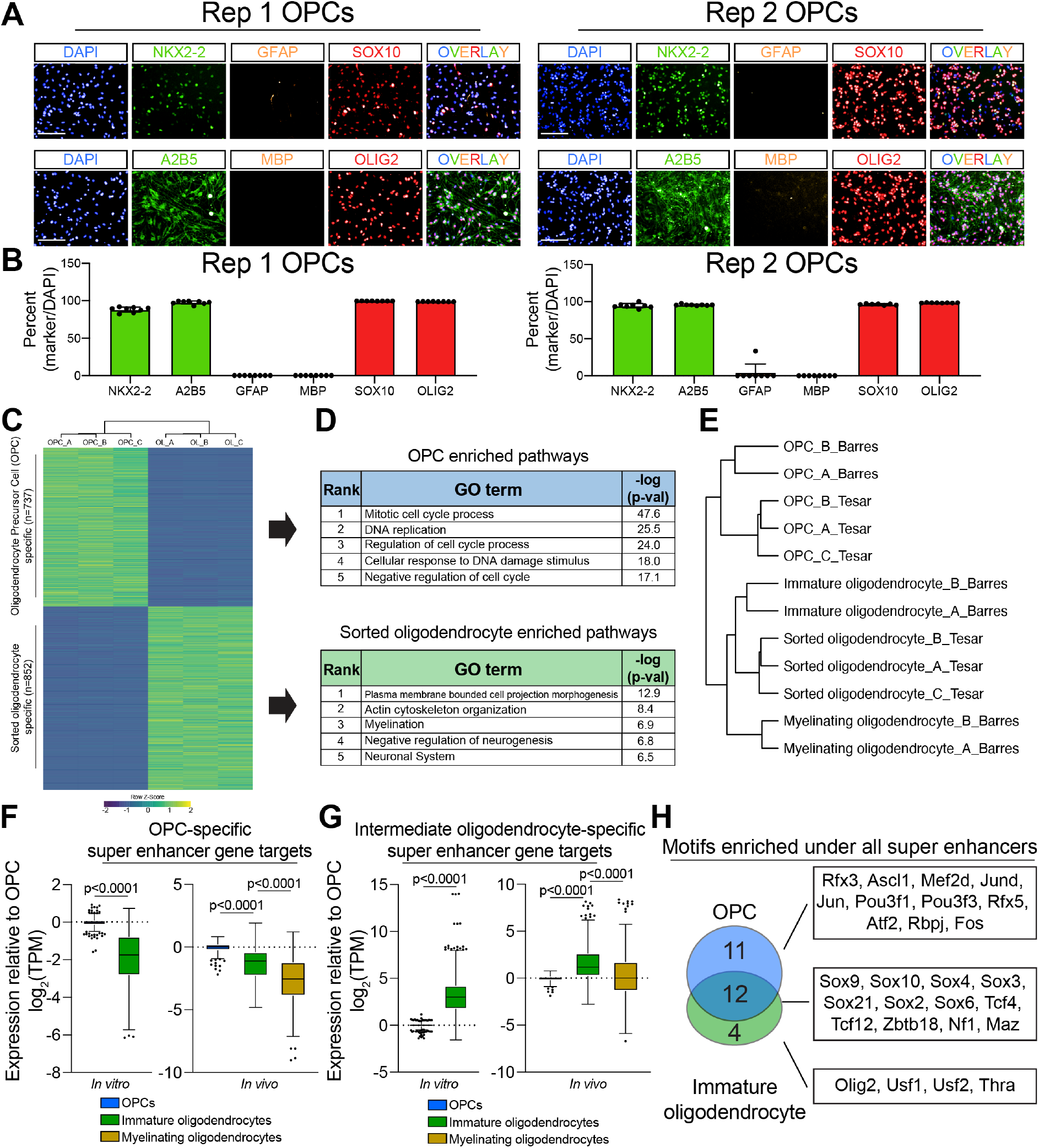
OPC and Immature Oligodendrocyte Super Enhancers are Enriched for Sox Family Motifs, Related to Figure 1. **(A)** Representative images of immunocytochemistry for OPC markers NKX2-2, OLIG2, A2B5, and SOX10 along with the oligodendrocyte marker MBP and astrocyte marker GFAP for both replicate 1 and replicate 2 OPCs. These replicates represent two independent batches of OPCs from different mouse strains. Nuclei are marked by DAPI (in blue). Scale bars, 100μm. **(B)** Quantification of OPC markers NKX2-2, OLIG2, A2B5, and SOX10, oligodendrocyte marker MBP, and astrocyte marker GFAP as a percent of total cells indicated by DAPI staining for both replicate 1 and replicate 2 OPCs. **(C)** Heatmap representation of significantly differentially expressed genes between OPCs and oligodendrocytes shown as row Z-score (log2FC >2, P-adj < 0.001). Columns were sorted by unsupervised hierarchical clustering and rows were ranked based on the fold change of gene expression in oligodendrocytes relative to OPCs. Each column represents an individual and independent RNA-seq sample using replicate 1 OPCs and magnetically purified oligodendrocytes. **(D)** Gene ontology (GO) analysis of genes that are significantly differentially expressed in OPCs (in blue) and oligodendrocytes (in green) (log2FC >2, P-adj < 0.001). Table shows the rank of the GO term along with –log(p-value). **(E)** Dendrogram of unsupervised hierarchical clustering of gene expression data from our *in vitro* OPCs and pre-myelinating oligodendrocytes (indicated by “Tesar”) and publicly available gene expression data from *in vivo* OPCs, immature oligodendrocytes, and myelinating oligodendrocytes datasets.^19^ **(F-G)** Box and whisker plot of change in gene expression (TPM) relative to OPCs for OPC-specific (F) and oligodendrocyte-specific (G) super-enhancer associated genes. Gene expression values were taken from our *in vitro* OPCs and immature oligodendrocytes and publicly available datasets for *in vivo* OPCs, immature oligodendrocytes, and myelinating oligodendrocytes.^19^ The black line represents the median with the box borders representing the upper and lower quartiles with dots representing statistical outliers. p-values were calculated using the Mann-Whitney test for *in vitro* plots and Kruskal Wallis One-Way ANOVA with Dunn's multiple comparisons test for *in vivo* plots. See also Table S1. **(H)** Venn diagram of the transcription factors expressed by the oligodendrocyte lineage (TPM >10) whose motifs were significantly enriched (p-value < 1×10^-10^) under superenhancers in OPCs (in blue) and immature oligodendrocytes (in green). See also Table S2.

**Figure S2.**
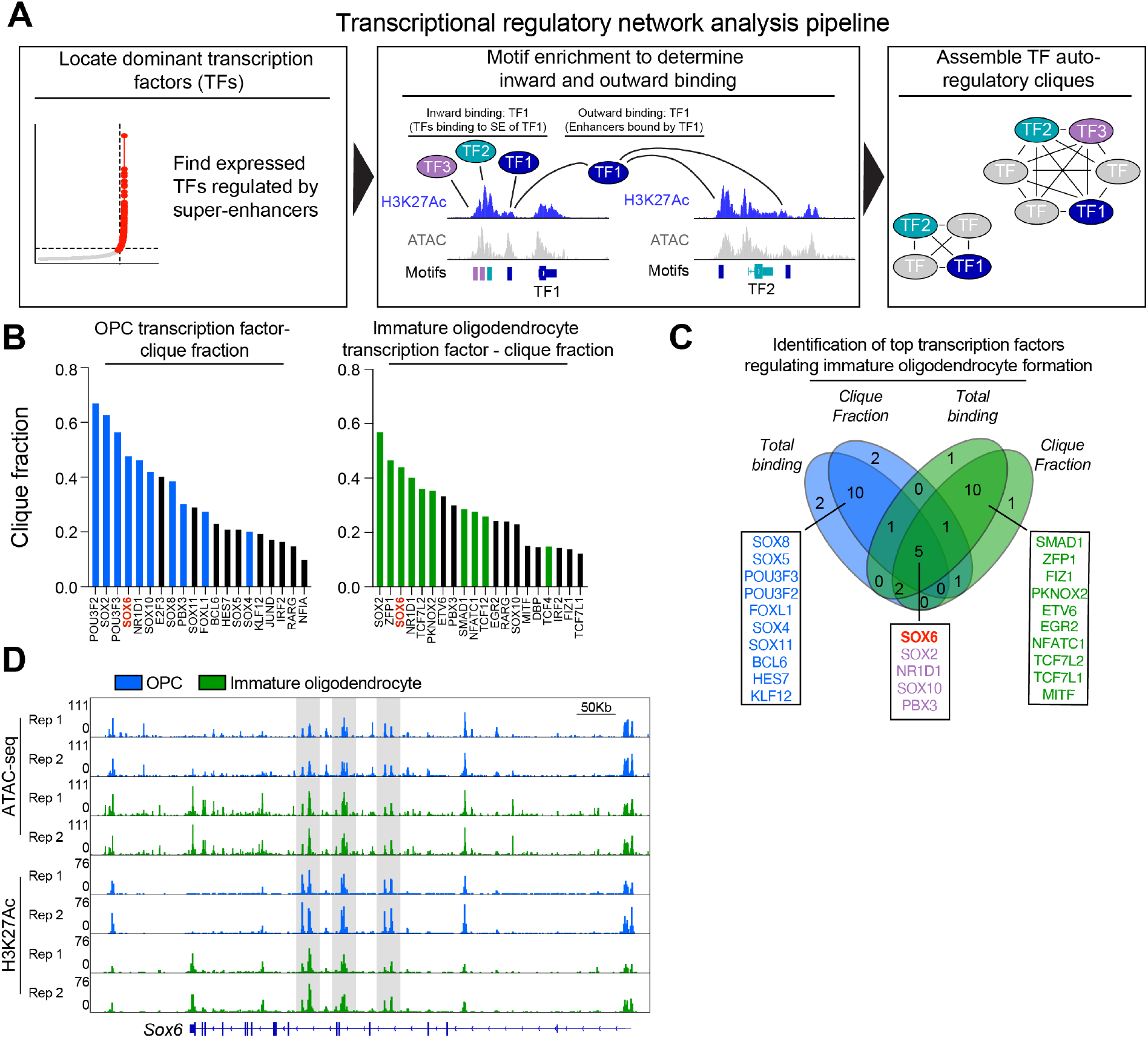
Transcriptional Regulatory Analysis Pinpoints SOX6 as a Dominant Transcriptional Regulator of Immature Oligodendrocyte Formation, Related to Figure 1. **(A)** Schematic representation of the transcriptional regulatory network analysis pipeline. First, expressed super-enhancer associated transcription factors are called based on ATAC-seq peaks within H3K27Ac defined super-enhancers. Then, the number of transcription factors that bind to open chromatin of a SE transcription factor loci is calculated (inward binding) and number of times a transcription factor binds to open chromatin at other SE transcription factor loci is calculated (outward binding). Lastly, the prevalence of a transcription factor in autoregulatory transcription factor circuits (or cliques) is calculated. **(B)** Bar graph of the top 20 super-enhancer associated transcription factors ranked by clique fraction, or their representation in transcriptional autoregulatory cliques, in OPCs (Left) and immature oligodendrocytes (right). Clique fraction equals the number of cliques that includes the transcription factor divided by the total number of cliques within the transcriptional regulatory network. Each bar represents a single transcription factor and the top 10 connected transcription factors are highlighted in blue (OPCs) and in green (immature oligodendrocytes). SOX6 is highlighted in red as the most highly connected transcription factor. See also Table S3. **(C)** Venn diagram overlapping the top 20 transcription factors in total binding and clique fraction in OPCs (in blue) and oligodendrocytes (in green). The top nodes specifically in OPCs and oligodendrocytes are listed in blue and green respectively while nodes shared between cell states are highlighted in purple. Genes are listed in order of their total binding. **(D)** Genome browser view of two replicates of ATAC-seq, and H3K27Ac ChIP-seq in OPCs (in blue) and oligodendrocytes (in green) at loci for *Sox6*. Super-enhancer loci are highlighted in gray. Scale bar, 50Kb.

**Figure S3.**
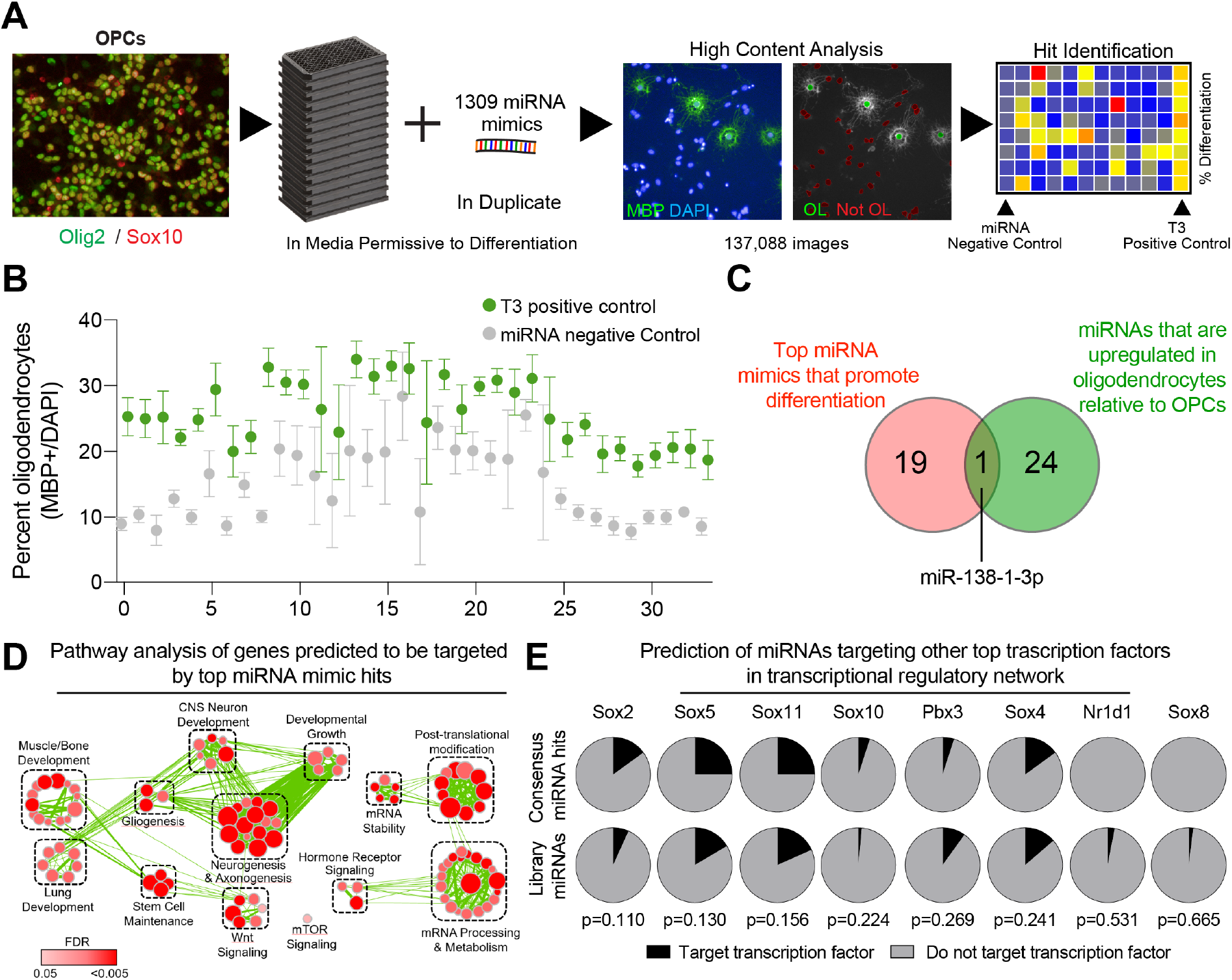
Phenotypic Screen Reveals That miRNAs That Drive Oligodendrocyte Formation Converge on Targeting SOX6, Related to Figure 2. **(A)** Schematic depicting the procedure for the primary miRNA mimic screen to uncover miRNAs that accelerate oligodendrocyte formation from OPCs. **(B)** Primary screen positive control (T3) and negative control (non-targeting miRNA mimic) percent oligodendrocyte (MBP+/DAPI) metrics on a per plate basis. Data represent mean ± SD from 8 technical replicates (individual wells) per plate. **(C)** Venn diagram indicating the overlap between top miRNA mimic hits from the primary screen that drive oligodendrocyte formation (in red) and miRNAs that significantly increase in oligodendrocytes relative to OPCs (in green). **(D)** Enrichment network of pathways of genes predicted to be targeted by top miRNA mimic hits. Each node represents a pathway and the size and color of each node is proportional to the number of genes and statistical significance of the pathway respectively. **(E)** Pie charts showing the number of miRNAs predicted to target other transcription factors (in black) to miRNAs that do not target that transcription factor (in gray) within top miRNA hits compared to their prevalence in the whole miRNA screening library. p-values were calculated using hypergeometric analysis.

**Figure S4.**
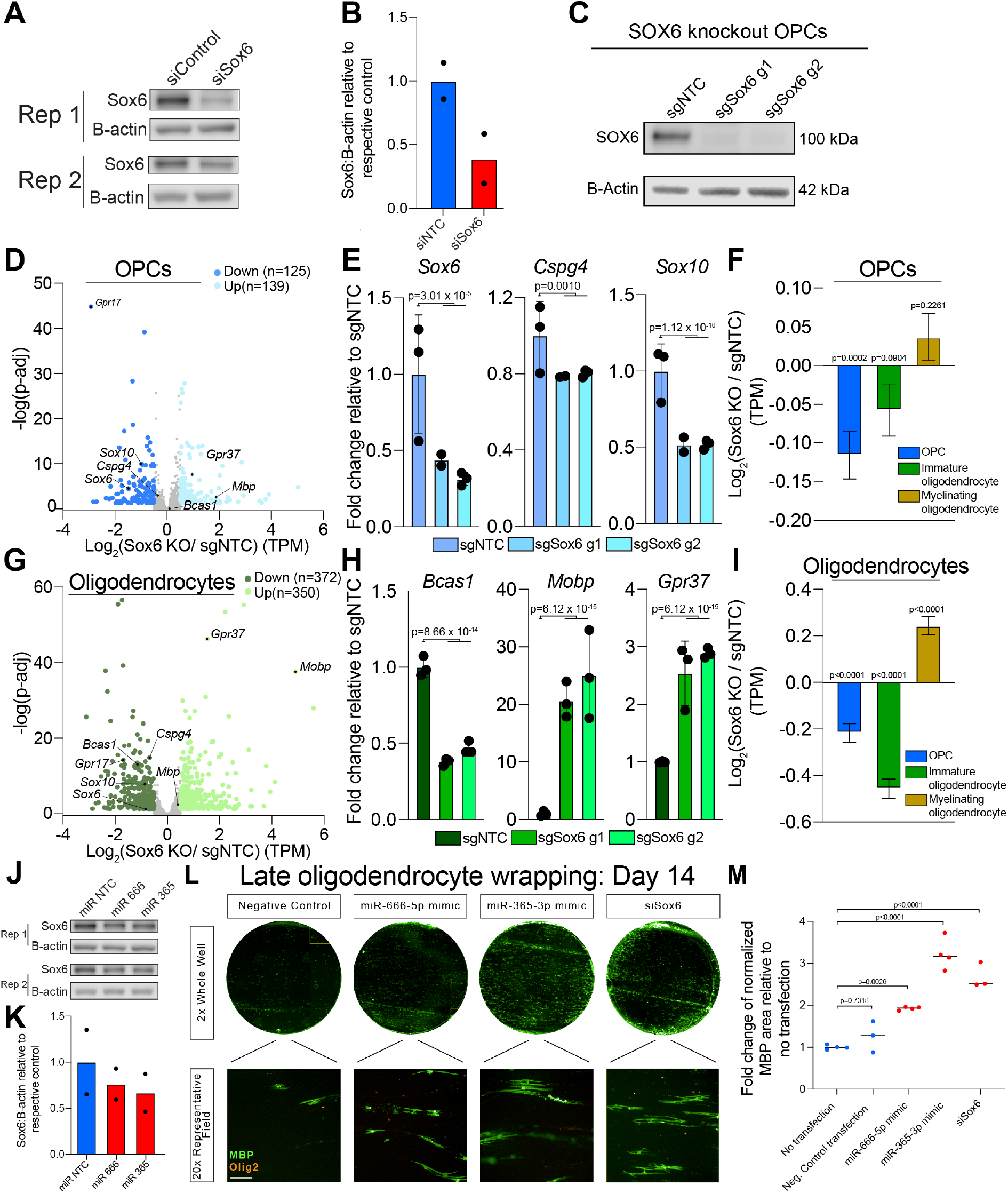
Loss of SOX6 Accelerates Oligodendrocyte Maturation, related to Figure 4. **(A)** Western blot of SOX6 from OPCs transfected with non-targeting (siNTC) or Sox6-targeting (siSox6) siRNAs. Data represent results using replicate 1 and replicate 2 OPCs. **(B)** Quantification of the ratio of SOX6 to B-actin of OPCs transfected with siRNA targeting SOX6 (in red) or non-targeting siRNA (siNTC in blue). Data are presented as the mean using replicate 1 and replicate 2 OPCs. **(C)** Western blot of SOX6 from nuclear lysates of control (sgNTC), and SOX6 knockout OPCs using two different guides (sgSox6 g1 and sgSox6 g2) with B-Actin as a loading control. **(D)** Volcano plot of differentially expressed genes (Log2FC greater than 0.5 or less than −0.5 and P-adj < 0.05) between control (sgNTC) and SOX6 knockout (combined sgSox6 g1 and sgSox6 g2) OPCs. Gray dots are genes not significantly different between conditions. Example genes from *in vivo* signature gene sets are labeled. Data are from 3 biological replicates per condition (independent samples), except for sgSox6 g1 OPCs, which has 2 biological replicates. **(E)** Quantification of the normalized number of transcripts (TPM) for *Sox6, Cspg4* and *Sox10* in control sgNTC OPCs (in blue), and SOX6 knockout OPCs sgSox6 g1 and g2. Data are normalized to sgNTC OPCs and represent mean ± SD from 3 biological replicates (independent samples) from RNA-seq, except for sgSox6 g1 OPCs, which has 2 biological replicates. **(F)** Quantification of normalized gene expression (TPM) of signature genes from *in vivo* OPCs, newly formed oligodendrocytes, and myelinating oligodendrocytes in SOX6 knockout OPCs relative to control OPCs (sgNTC). Data are presented as mean ± SEM and p-values were calculated using the one-sample Wilcoxon signed rank test. **(G)** Volcano plot of differentially expressed genes (Log2FC greater than 0.5 or less than −0.5 and P-adj < 0.05) between control (sgNTC) and SOX6 knockout (combined sgSox6 g1 and sgSox6 g2) oligodendrocytes at day 3 of differentiation. Gray dots are genes not significantly different between conditions. Example genes from *in vivo* signature gene sets are labeled. Data are from 3 biological replicates per condition (independent samples). **(H)** Quantification of the normalized number of transcripts (TPM) of intermediate oligodendrocyte gene *Bcas1* and myelinating oligodendrocyte genes *Gpr37* and *Mobp* in control sgNTC oligodendrocytes (in green), and *Sox6* knockout (sgSox6 g1 and g2) oligodendrocytes. Data are normalized to sgNTC oligodendrocytes and represent mean ± SD from 3 biological replicates (independent samples) from RNA-seq, except for sgSox6 g1 OPCs, which has 2 biological replicates. **(I)** Quantification of normalized gene expression (TPM) of signature genes from *in vivo* OPCs, newly formed oligodendrocytes, and myelinating oligodendrocytes in *Sox6* knockout oligodendrocytes relative to control oligodendrocytes (sgNTC). Data are presented as mean ± SEM and p-values were calculated using the one-sample Wilcoxon signed rank test. **(J)** Western blot of SOX6 from OPCs transfected with non-targeting miRNA (miR NTC) or putative SOX6-targeting miRNAs miR-666 and miR-365. Data represent results using replicate 1 and replicate 2 OPCs. **(K)** Quantification of the ratio of *Sox6* mRNA to *B-actin* of OPCs transfected with miR-666, miR-365, or non-targeting miRNA (miRNA NTC in blue). Data are presented as the mean using replicate 1 and replicate 2 OPCs. **(L)** Representative immunocytochemistry images of myelinating oligodendrocytes (MBP+ in green) on microfibers with all cells of the oligodendrocyte lineage labeled with OLIG2 (in orange). Scale bar, 100μm. **(M)** Quantification of the area of MBP+ myelinating oligodendrocytes normalized to the number of OLIG2+ cells per image. Data are normalized to the no transfection condition and presented as mean ± SD from 3 separate microfiber well inserts per condition.

**Figure S5.**
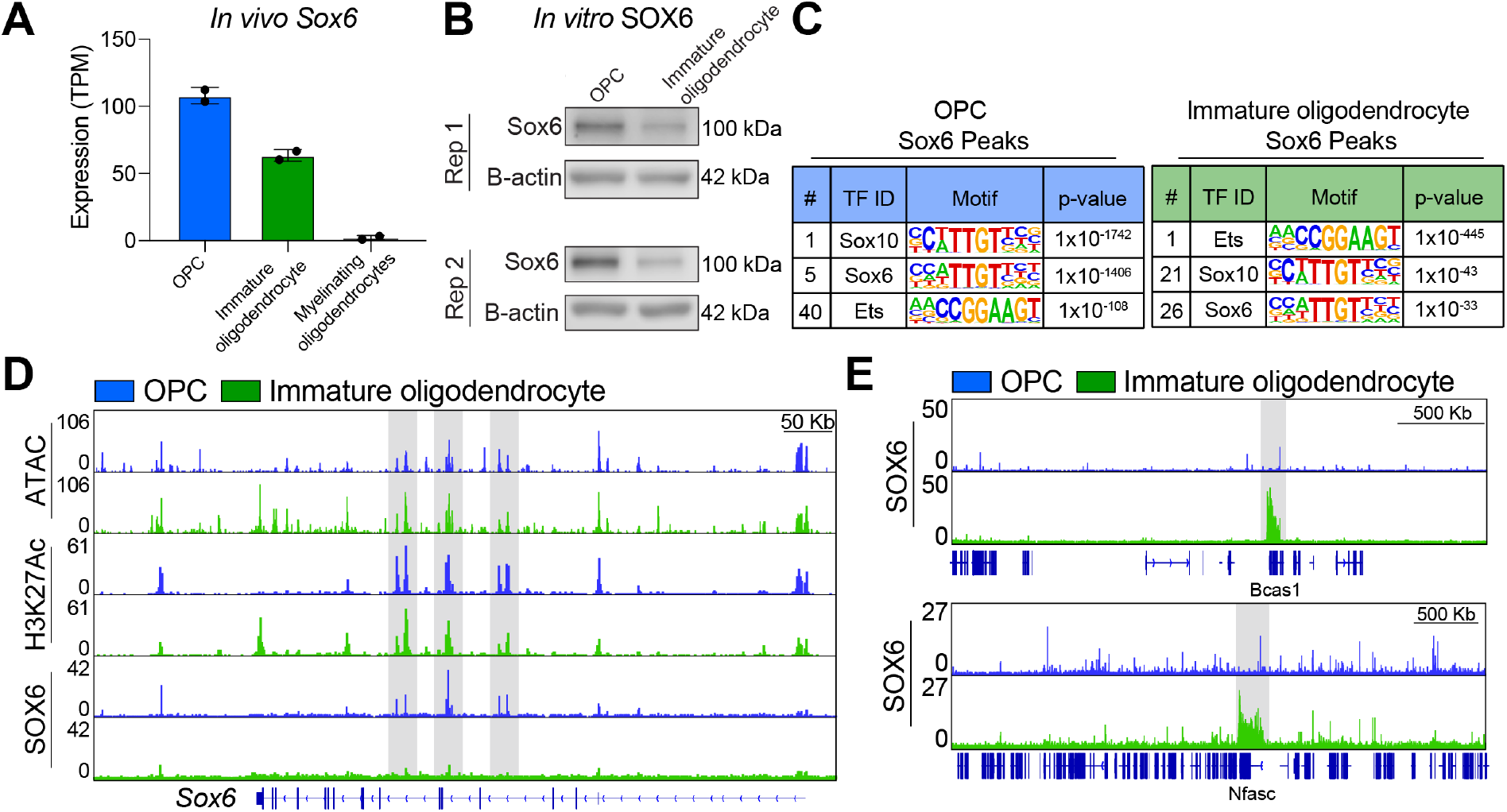
SOX6 Reorganizes from OPC Super Enhancers to Form Clusters Across Gene Bodies in Immature Oligodendrocytes, Related to Figure 4. **(A)** Quantification of the normalized number of transcripts (TPM) for *Sox6 in vivo* OPCs (in blue), immature oligodendrocytes (in green), and myelinating oligodendrocytes (in orange). Data represent mean ± SD from 2 biological replicates (independent samples) from publicly available RNA-seq. **(B)** Western blot of SOX6 from nuclear lysates of OPCs compared to sorted immature oligodendrocytes with B-Actin as a loading control using replicate 1 and replicate 2 OPCs. **(C)** Table of known motifs significantly enriched under SOX6 peaks in OPCs (in blue) and immature oligodendrocytes (in green). Charts display the transcription factor name, motif, and p-value ranked in order of significance (# indicates rank out of all 1006 motifs in the analysis). See also Table S2 for full list of ranked motifs for each cell type. **(D)** Genome browser view of ATAC-seq, H3K27Ac ChIP-seq, and SOX6 ChIP-seq in OPCs (in blue) and oligodendrocytes (in green) at *Sox6*. Super-enhancer loci are highlighted in gray. Scale bar, 50 Kb. **(E)** Zoomed out genome browser view of SOX6 ChIP-seq in OPCs (in blue) and oligodendrocytes (in green) at example SOX6 condensate loci including *Bcas1* and *Nfasc*. SOX6 condensates are highlighted in gray. Scale bars, 500Kb.

**Figure S6.**
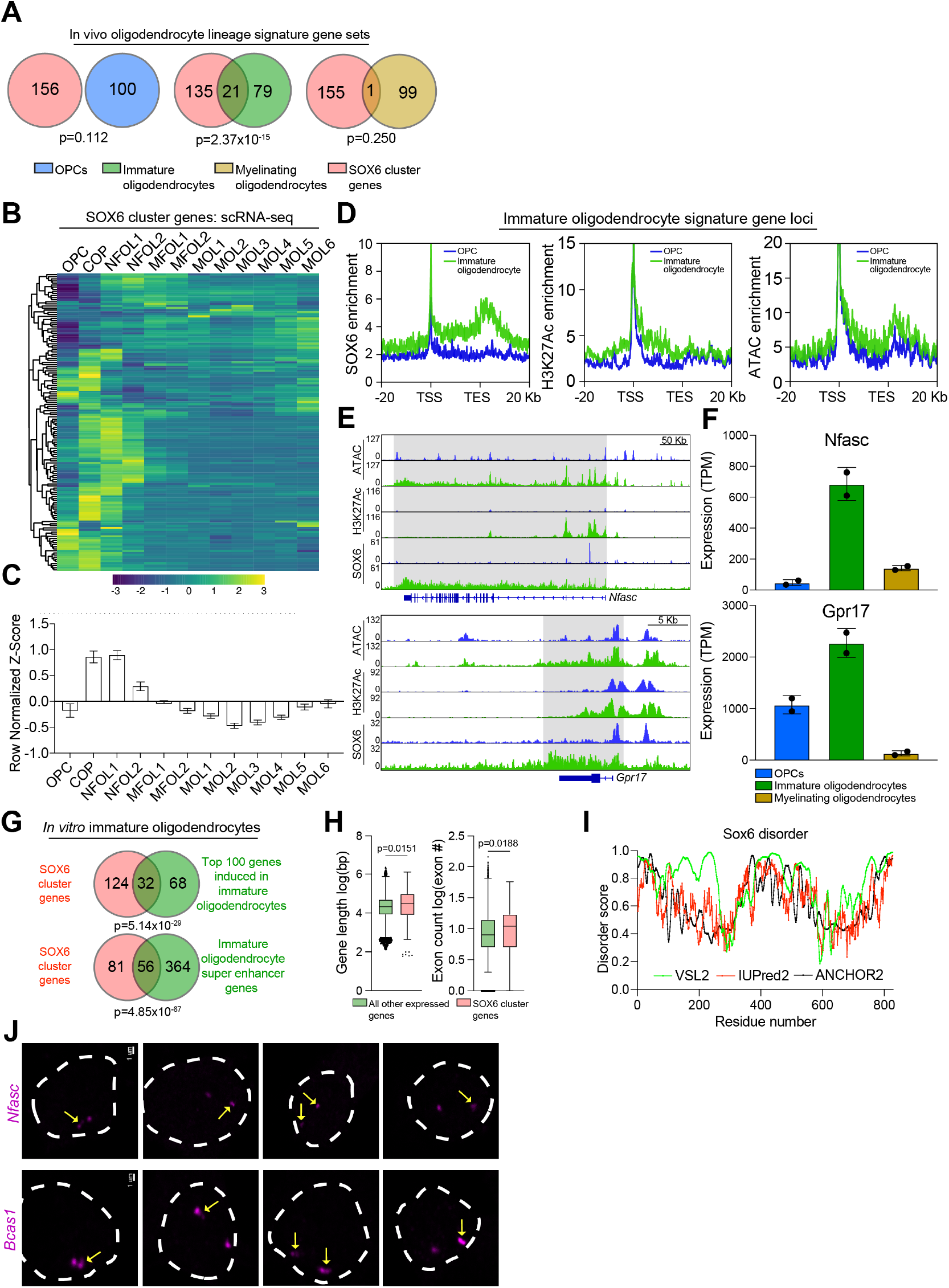
SOX6 at Decondensed Genes Transiently Expressed During Oligodendrocyte Differentiation, Related to Figure 5. **(A)** Venn diagram indicating the overlap between SOX6 cluster genes (in red) and the top 100 signature genes in *in vivo* OPCs (in blue), immature oligodendrocytes (in green), and myelinating oligodendrocytes (in orange). p-values were calculated by hypergeometric analysis. See also Table S5 for full list of *in vivo* signature genes. **(B)** Heatmap representation of row normalized expression of SOX6 cluster genes (log[average UMI]) from single-cell RNA-seq of the oligodendrocyte lineage in the developing mouse brain. COP = committed oligodendrocyte progenitor, NFOL = newly formed oligodendrocyte, MFOL = myelin forming oligodendrocyte, and MOL = mature oligodendrocytes. **(C)** Bar graph representation of the normalized Z-scores of the SOX6 condensate genes for each cell cluster from the single-cell RNA-seq of the oligodendrocyte lineage in the developing mouse brain. Abbreviations are the same as in Figure S5C. **(D)** Aggregate plots of ATAC-seq, H3K27-Ac ChIP-seq, and SOX6 ChIP-seq across the gene bodies of immature oligodendrocyte signature genes and 20Kb upstream from the transcription start site (TSS) and 20Kb downstream from the transcription end site (TES). **(E)** Genome browser view of ATAC-seq, H3K27Ac ChIP-seq, and SOX6 ChIP-seq in OPCs (in blue) and pre-myelinating oligodendrocytes (in green) at example SOX6 condensate loci including *Fyn*, and *Gpr17*. SOX6 clusters are highlighted in gray. Scale bars, 50Kb and 5Kb respectively. **(F)** Quantification of the normalized number of transcripts (TPM) for *Fyn*, and *Gpr17* from *in vivo* OPCs (in blue), immature oligodendrocytes (in green), and myelinating oligodendrocytes (in orange). Data represent mean ± SD from 2 biological replicates (independent samples) from publicly available RNA-seq. **(G)** Venn diagram indicating the overlap between SOX6 cluster genes (in red) and the top 100 genes induced in oligodendrocytes relative to OPCs (top, in green) and separately with immature oligodendrocyte super-enhancer associated genes (bottom, in green). p-values were calculated by hypergeometric analysis. **(H)** Box and whisker plot of gene length (log (bp)) and number of exons in all expressed genes in immature oligodendrocytes and SOX6 cluster genes. The black line represents the median with the box borders representing the upper and lower quartiles with dots representing statistical outliers. p-values were calculated using the Mann Whitney test. **(I)** Disorder analysis of SOX6 (UniProt: P40645). The algorithms used were: VSL2 (in green), IUPred2 (in red), and ANCHOR2 (in black). An amino acid score above the dotted line indicates a disordered score greater than 0.5 and that the amino acid sequence is disordered. Sequence is written from N-terminus to C-terminus. **(J)** Representative images of nascent RNA FISH of *Bcas1* and *Nfasc* (magenta) in fixed immature oligodendrocytes representing gene melting. Yellow arrows indicate elongated gene melting events.

## SUPPLEMENTAL TABLES (provided as separate .xlsx files)

**Table S1. Super enhancer associated genes and miRNAs**

List of expressed super enhancer associated genes and miRNAs in OPCs and immature oligodendrocytes and raw dynamic enhancer outputs for both replicates and cell states.

**Table S2. HOMER motif enrichment analysis results**

HOMER motif analysis was performed under ATAC-seq peaks (FDR < 0.001) that intersected OPC and immature oligodendrocyte super enhancers. HOMER motif analysis was also performed under SOX6 peaks (FDR<0.001) in OPCs and immature oligodendrocytes.

**Table S3. Transcriptional network analysis**

Transcriptional network analysis was used to elucidate the connectedness (inward and outward binding) and prevalence in self-reinforcing auto-regulatory cliques of superenhancer associated transcription factors in OPCs and immature oligodendrocytes.

**Table S4. miRNA phenotypic screen results**

Primary miRNA screening data showing the impact of 1,295 miRNA mimics in duplicate on percentage of oligodendrocytes (MBP+/DAPI) relative to T3 baseline controls. miRNA top hits were called using a combinatorial score based on percent oligodendrocytes relative to positive and negative controls on each plate and are highlighted in the table.

**Table S5. *In vivo* state-specific signature genes**

List of the top 100 genes that are most strongly enriched in OPCs, immature oligodendrocytes, or myelinating oligodendrocytes compared to the other two states using publicly available *in vivo* RNA-seq data.

**Table S6. SOX6 cluster genes**

List of expressed genes that harbor SOX6 intensity across their gene bodies in immature oligodendrocytes that is greater than 2-fold over background (input) and greater than 2-fold over SOX6 intensity in OPCs. Metascape pathway analysis results are also included as a separate sheet.

